# Targeting *Leishmania donovani* Sphingosine Kinase 1 using PF-543 enhances immune response and limits parasite load

**DOI:** 10.1101/2025.05.05.652326

**Authors:** Evanka Madan, Ruby Bansal, Jhalak Singhal, Monica Saini, Sadat Shafi, Nishant Joshi, Shailja Singh

## Abstract

**Background:** Sphingosine-1-phosphate (S1P) is a bioactive lipid mediator regulating apoptosis, proliferation, and immune responses. While S1Ps presence in *Leishmania donovani* phagolysosomes has been reported, the role of sphingosine kinases, especially SphK1, in parasite survival and host immune modulation remains underexplored. This study investigates the molecular and functional role of *L. donovani* SphK1 (*Ld*SphK1) and evaluates the antileishmanial potential of PF-543, a specific SphK1 inhibitor.

**Methods:** *Ld*SphK1 and human SphK1 (*rh*SphK1) were cloned, expressed in *E. coli*, purified, and analyzed by SDS-PAGE. Enzymatic activity and inhibition by PF543 were assessed using NBD-S1P-based fluorometric assays. Protein-ligand interactions were analyzed using Microscale Thermophoresis (MST). *Leishmania* promastigotes overexpressing *Ld*SphK1 were studied via confocal microscopy, and their viability and infectivity were assessed *in vitro*. THP-1 macrophages infected with *L. donovani* were treated with PF543 alone or with Amphotericin B and analyzed by MTT assay, RT-PCR, Giemsa staining, ELISA and immunoblotting. *In vivo* efficacy was tested in *L. donovani*-infected Swiss mice.

**Results:** *rLd*SphK1 (∼102 kDa) and *rh*SphK1 (∼50 kDa) were enzymatically active and significantly inhibited by PF-543. MST confirmed high-affinity binding of PF-543 (KD ∼29 microMolar). In *L. donovani* SphK1 overexpressor *(Ld*SphKa) promastigotes, PF543 inhibited SphK1 activity and reduced parasite infectivity, more than in wildtype *L. donovani* promastigotes. Notably, PF543 treatment reduced parasite infectivity *in vitro*, lowered amastigote load by ∼40%, and promoted a pro-inflammatory cytokine shift (↑IL-12, ↑TNF-α, ↓IL-10). Inhibition of ceramide synthesis and S1P supplementation revealed that S1P rescues ceramide-induced parasite death, implicating SphK1 in parasite survival. PF543 and Amphotericin B demonstrated synergistic anti-parasitic effects both *in vitro* and *in vivo*, with >90% reduction in parasite burden in mice.

**Conclusion:** PF543 is a potent inhibitor of SphK1, impairing parasite survival and modulating host immune responses. When combined with Amphotericin B, it offers a synergistic therapeutic strategy against visceral leishmaniasis, warranting further clinical exploration.

**Author Summary:** Leishmaniasis, a neglected tropical disease, has limited available treatments and is becoming more resistant to medications. In this study, we explored the therapeutic potential of PF-543, a potent sphingosine kinase 1 (SphK1) inhibitor (demonstrated anticancer effects in various preclinical models) against *Leishmania donovani*. We successfully cloned and purified both *Leishmania* and human SphK1 proteins and confirmed PF-543 binding through biochemical and biophysical assays. Overexpression of *Ld*SphK1 in parasites enhanced their survival and infectivity. *In vitro,* PF-543 treatment of infected macrophages decreased amastigote burden, shifted cytokine profiles towards a pro-inflammatory state, and enhanced host cell apoptosis. Notably, PF-543 acted synergistically with Amphotericin B, the current clinical drug, both *in vitro* and *in vivo* in Swiss mice, drastically lowering parasite burden. This study highlights the possibility of combination therapy and finds PF-543 to be a promising irresistible host-targeted antileishmanial drug.

**Graphical Abstract:** 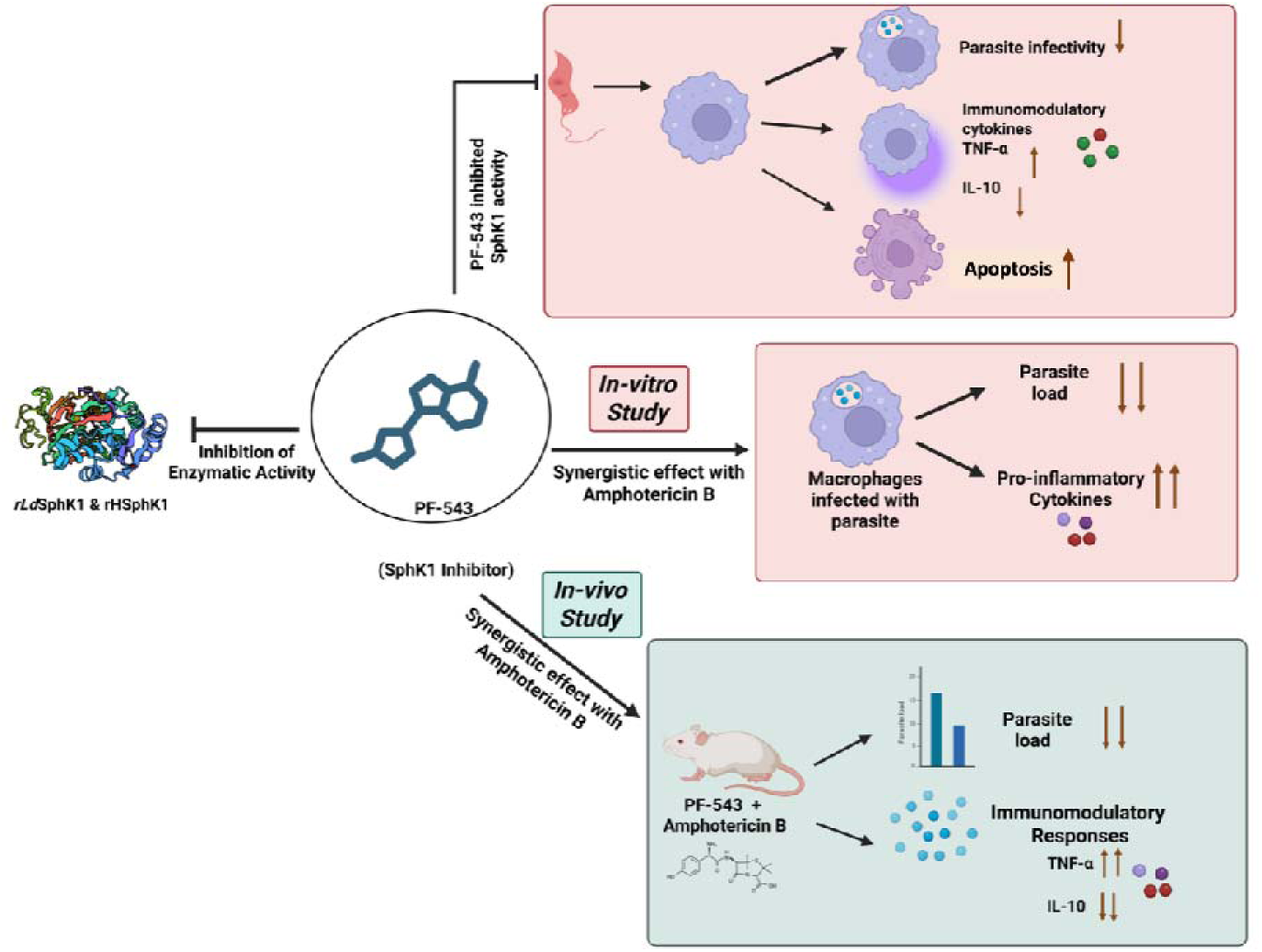

## Introduction

Leishmaniasis is a neglected tropical disease caused by intracellular protozoa parasite *Leishmania* [1]. It invades macrophages and selectively impairs host’s critical signaling pathways for its successful intracellular growth and proliferation [2]. In particular, alteration in lipid metabolic pathways, lipid relocation, accumulation and mitochondrial dysfunction during *Leishmania* infection has been proven to be a critical step in the progression of disease [3].

Decrease in efficiency of current anti-leishmanial agents, increase in cost and complexity of curing *Leishmania*, increased parasitic resistance to the chemicals and limited number of available anti-leishmanial drugs has spur the research for novel approaches [4] Pentavalent antimonials (sodium stibogluconate and meglumine antimoniate), liposomal amphotericin B, Miltefosine and Paromomycin are the current drugs for Visceral Leishmaniasis [5,6]. However, resistance has been seen in endemic regions, where these drugs are extensively used, with risk of serious cardiotoxicity, toxic effects on the kidneys, leading to death [6,7]. The need for novel, effective, and safer therapeutic alternatives is therefore urgent.

In this context, targeting host or parasite-encoded enzymes has emerged as the promising strategy for identifying new anti-leishmanial agents. For the first time, we explored the potential of PF-543, a potent Sphingosine kinase (SphK) inhibitor to modulate Sphingosine-1-phosphate (S1P) signalling, and disrupt *Leishmania* infection. Interestingly, evidences have suggested a key role for SphK in several lung pathologies, including lung fibrosis, pulmonary artery hypertension, asthma, ischemia reperfusion injury [8–12]. Recent studies have also implicated S1P signaling in the biology of *Leishmania donovani,* especially within phagolysosomes [13].

Among the two SphK isoforms, SphK1 is more responsive to various stimuli and is increasingly viewed as a favourable therapeutic target compared to the more constitutively active SphK2 [14]. Our work focused on modulating SphK1, a key enzyme in conversion of sphingosine to SIP, which is critical for parasite survival and host immune evasion. Notably, SphK1 enables *Leishmania* to manage toxic metabolites, endure cellular stress, and establish infection, making it a compelling drug target [15].

These days, there are growing interests in using small-molecule inhibitors of the Sphingosine kinases (SphK1 and SphK2) as potential therapeutics or preventive agents for human diseases [16]. The SphK inhibitors: N, N-dimethyl-sphingosine (DMS), SKI-5C and PF-543 are formerly developed as antitumoral drugs [17]. We hypothesize that targeting both host and parasite SphK1 with these inhibitors, especially PF-543, could suppress parasite survival within macrophages and may serve as a novel therapeutic approach to the classical treatments against *Leishmania*.

Till date, there is no study addressing the role of PF-543 in sensing SIP signalling and inducing lysosomal stress in human macrophages infected with *Leishmania*, so, we sought to study the underlying molecular pathways of these SphK inhibitors involved in parasite survival, including the role of ceramide accumulation and intracellular calcium dysregulation, which are known to contribute to parasite death via ER stress and autophagy [18–21].

Notably, both *Leishmania* and macrophages are known to harbor SphK1, making PF-543 a unique dual-target inhibitor. Importantly, PF-543-mediated inhibition of host SphK1 appears safe due to the compensatory presence of SphK2, further minimizing potential side effects and reducing the likelihood of resistance—since the drug targets a host rather than a parasite protein.

In this study, we cloned and expressed *Leishmania* and human SphK1, confirmed protein purification, and validated PF-543 binding through Microscale Thermophoresis (MST), *in silico* docking, and kinetic analysis. In *Leishmania* promastigotes, SphK1 was overexpressed and checked for SphK1 activity, parasite infectivity, inflammatory responses following PF-543 treatment. Mechanistic studies explored the role of ceramide and autophagy, while potential impact of Amphotericin B, the current drug used in clinics alone and in combination with PF-543 against *L. donovani* promastigotes, amastigotes and Swiss mice was evaluated.

Usage of Amphotericin B liposome as anti-leishmanial is already published [22] and patented [23]. PF-543, the most potent SphK1 inhibitor developed by Pfizer, is not yet known to be an anti-leishmanial drug. Here, for the first time, we report the anti-leishmanial potential of PF-543, both as standalone agent and in combination with Amphotericin B as a new anti-leishmanial formulation.

By leveraging host-targeted mechanisms and minimizing the risk of drug resistance, the PF-543 and Amphotericin B combination introduces a promising new direction in leishmaniasis treatment. These findings not only provide insight into host-parasite interactions but also open up new avenues for exploiting host pathways to combat parasitic diseases.

## Methods

### In silico search for SphK1 like protein in Leishmania donovani

An *in-silico* search was conducted in the Kinetoplastid database TriTrypDB (www.tritrypdb.org) to identify any homologous protein similar to prokaryotic SphK1 in *Leishmania donovani* [24]. Sequence alignments and identity computations were conducted using ClustalW [25]. Homologues of SphK1 from *L. donovani*, along with other prokaryotes and eukaryotes were analyzed using the Molecular Evolutionary Genetics Analysis (MEGAX) tool [26]. The evolutionary trajectory was determined using the Neighbor-Joining method. The bootstrap consensus tree was generated from replicates and the evolutionary distances were calculated using the Poisson correction method. Pymol was utilized to align the two structures and compute the RMSD post-alignment [27].

### In-silico docking studies

The protein structure of hSphK1 (Uniprot ID-Q9NYA1), a protein derived from H. sapiens, was retrieved from the protein data bank [28,29]. Protein structure of *L. donovani* sequence *Ld*SphK (*Ld*SphK_260680.1.1) was obtained from TriTrypDB [30]. Phyre2 web server was used for homology 3D protein modelling of the protein and Procheck sever was used for Ramachandran plot analysis of the model [31,32]. The Swiss PDB viewer and ChemBio Draw ultra 3D software were used to optimize the structure of proteins and ligands [33,34]. The process of molecular docking between molecule PF-543 and hSphK1 was conducted using Autodock version 4.2 and Cygwin terminal version 3.1 software [35,36]. The residues located in the vicinity of the binding pocket were selected to create a grid (with a spacing of 0.375) centred at coordinates X-51.488, Y-53.406, Z--1.49 for hSphK1 and X-4.021, Y- - 21.535, Z- 15.254 for *Ld*SphK1 respectively. This grid was used to facilitate the binding of the ligand. The docking data were further analyzed and visualized using the PLIP, Ligplot+ version 2.2, Discovery Studio version 19.1.0, and Pymol version 2.3.2 software tools [37–40].

### Cloning and recombinant protein purification of LdSphk1

Nucleotide sequence encoding full length sequence of *L. donovani* Sphk1 (1-2808 bp, **Fig S1A**) was selected for recombinant protein generation as a fusion protein with an N-terminal 6x- Histidine-tag using the vector pET-28 a(+) vector. DNA fragment for the gene cloning was PCR amplified from genomic DNA of *L. donovani* using the following primer pair: *Ld*Sphk1_BamHI_FP 5’-GATGGGATCCATGCACTCTTTCACCTCCAA-3’ and *Ld*Sphk1_XhoI_RP 5’-ATTCTCGAGCTACCCACTGCGCACGAGCT-3’with Phusion™ High-Fidelity DNA Polymerase (Thermo Scientific, US). The amplified DNA fragment was purified with QIAquick Gel Extraction Kit (Qiagen) using the manufacturer’s protocol.

Purified *Ld*Sphk1 insert and the expression vector pET-28 a(+) were digested with BamHI/XhoI restriction enzymes (New England Biolabs, UK), and were ligated overnight at 16°C using T4 DNA ligase (New England Biolabs, UK). The ligation mix was transformed into *E. coli* DH5-α competent cells (*E.Coli* DH5-α cells were obtained from ThermoFisher Scientific, USA Catalog number C0112) and positive clones were screened by colony PCR followed by confirmation of the cloned plasmid with BamHI/XhoI restriction digestion **(Fig S1B)** and further transformed in Rossetta *E. coli* (Rossetta *E. coli* were obtained from Sigma-Aldrich, USA with Catalog number 71397) expression strain for *Ld*Sphk1 recombinant protein expression **(Fig S1C).** For recombinant protein induction, the cells were induced with 1mM IPTG for 4 hours at 37°C in Terrific broth. Purification of recombinant protein was achieved by affinity chromatography.

The *rLd*SphK1 protein (amino acid sequence for the protein shown in **Fig S2A**) was eluted after addition of 150mM and 250mM imidazole solution. The eluted fractions were analysed by running on 12% SDS PAGE and the fractions containing pure protein were pooled and concentrated using a centricon device, followed by buffer exchange with PBS. Since most of the protein was localized in the inclusion bodies, we purified the protein from inclusion bodies under denaturation conditions. Briefly, inclusion bodies were obtained after sonication and subsequent centrifugation at 13,000 rpm and solubilized in urea buffer composed of 8M urea, 20mM Tris, 250mM NaCl with pH 8.0 and subsequently incubated with Ni-NTA beads (Qiagen, Germany) overnight for binding. The beads were then loaded in gravity flow column (Qiagen, Germany). Further, *Ld*SphK1 protein bound with the beads was then eluted by 10mM, 25mM, 50mM, 100mM, 250mM and 500mM imidazole solutions prepared in urea buffer. Eluted fractions containing pure protein were pooled and refolded by dialyzing against refolding buffer composed of 100mM Tris, 20% Glycerol, 250mM L-Arg, 1mM EDTA, 1mM GSG and 0.5mM GSSG with pH 8.0. Refolded protein was reconstituted in PBS (pH 7.4) and concentrated using centricon tubes with 10kDa cut-off (Merck, Germany).

### Recombinant protein purification of HumanSphk1

CDS encoding full-length SphK-1 (1155 bp) (Gene ID-8877) was amplified using the following primers: NheI_FP: 5′-AATGCTAGCATGGATCCAGCGGGCGGCCCC-3′ and XhoI_RP: 5′-TGGCTCGAGCTATAAGGGCTCTTCTGGCGGTG-3′.

The amplified DNA fragment was cloned between NheI and XhoI restriction sites of the pET-28a (+) expression vector, and the recombinant plasmid was transformed into *E. coli* strain BL21 (LDE3) gold. The *rhSphK1* protein was eluted after addition of 100mM, 250mM and 500mM imidazole solution **(Fig S2B).** Overexpression of 6×His-SphK-1 (*rSphK-1*) was induced with 1mM IPTG (Sigma-Aldrich) at an optical density (OD600) of 0.6, for 4 h at 37°C. The protein was purified using Ni-NTA agarose resin (Qiagen). Briefly, bacterial cells (Rosetta host strains DE3 competent cells (Derivatives of BL21) (Rossetta *E. coli* were obtained from Sigma-Aldrich, USA with catalog number 71397) were used to enhance the production of *rh*SphK-1 by using codons that are infrequently used in *E. Coli*) were harvested (6000L×Lg for 15 min) and lysed by sonication in a lysis buffer containing 50mM Na_2_HPO_4_ (pH 7.4) and 200mM NaCl, supplemented with 500μg/mL Lysozyme (Sigma-Aldrich), and 1mM Phenylmethylsulfonyl Fluoride (PMSF; ThermoFisher Scientific). The bacterial lysate was centrifuged at 10,000L×Lg for 20 min, and the supernatant obtained was loaded onto a Nickel-Nitrilotriacetic Acid (Ni-NTA) agarose resin (Qiagen)-packed column along with 10 mM imidazole, for 4h at 4°C with mild agitation. The protein was eluted with a continuous imidazole gradient of 25mM, 50mM, 100mM, 250mM, and 500mM. The protein purification was validated by 12% SDS-PAGE, followed by immunoblotting with anti-His tag antibody **(Fig S2B)** (raised in Swiss mice (obtained from the Central Laboratory Animal Resources, Jawaharlal Nehru University, New Delhi) using standard protocol approved by IAEC, JNU).

### Microscale thermophoresis (MST)

To validate the biophysical interaction between *rLd*SphK1 or *rHuman* SphK1 with PF-543 and analyze the binding and kinetics, Microscale Thermophoresis (MST) was performed in NanoTemper Monolith NT.115 instrument. Briefly, 20μM of *rLd*SphK1 or *rHuman* SphK1 in HEPES-NaCl buffer, pH 7.4 was labeled with 30μM Lysine reactive dye (5ul diluted in 200ul *rLd*SphK1 or *rHuman* SphK1 protein) using NanoTemper’s Protein Labelling Kit RED-NHS (L001, NanoTemper technologies, Germany) and incubated in the dark for 30 minutes at RT. Following incubation, the labelled *rLd*SphK1 or *rHuman* SphK1 protein along with the buffer was passed through an equilibrated column (provided in the kit). Fractions of the labelled protein were eluted followed by fluorescence count. Fluorescence counts ranging between 250-450 were taken. 5µM PF-543, diluted in HEPES/0.05% Tween-20, was titrated against the constant concentration of the labelled *rLd*SphK1 or *rHuman* SphK1 protein. Pre-mixed samples were incubated for 15 minutes before centrifugation at 8000 rpm for 10 minutes. The samples were filled into the capillaries treated with standards (K002 Monolith NT.115) and thermophoretic mobility was determined. All experiments were performed at RT, at 40% MST power and 20% LED power. Data evaluation was performed with the Monolith software (Nano Temper, Munich, German).

### Biochemical Characterization of Purified rLdSphK and rHumanSphK

The inhibition of *rLd*SphK1 catalytic activity was evaluated in a dose dependent manner with PF-543’s concentration ranging from 1 to 5μM. To evaluate the inhibitory effect of PF-543 on the enzymatic activity of *rLd*SphK-1, the protein was pre-incubated with different concentrations of PF-543 (0 to 5μM) for 45 min., followed by initiating the reaction in a buffer containing 0.05% Triton X-100, 1mM ATP, 2% DMSO and NBD-Sph (10μM). The fluorescence intensity of NBD-S1P was measured as described above. The results represented a maximum inhibition at 5μM of PF-543, while the protein concentration was kept constant at 200ng.

To detect K_m_ and V_max_ values, varying concentration of NBD Sphingosine (2-20μM) was taken along with 200ng of r*Ld*SphK1. For this, the recombinant protein was incubated with 200μM ATP, NBD–Sphingosine (NBD-Sph) was used as a substrate and conversion of NBD-Sph to NBD-S1P was evaluated. Readings were taken real-time after every 5 sec for 15-20 min at an excitation/emission wavelength of 490/530 nm. Standard curve of SIP was plotted. The enzymatic activities were calculated by plotting fluorescence/time vs substrate concentration followed by plotting of fluorescence/pmoles (velocity) vs substrate concentration. Finally 1/Vmax vs 1/[S] was plotted to calculate Km and Vmax according to Lineweaver-Burk plots at 37°C (Lineweaver and Burk, Science Direct, 2011).

### Leishmania cell culture

*Leishmania donovani Bob* promastigotes (*Ld*Bob strain/MHOM/SD/62/1SCL2D, originally obtained from Dr Stephen Beverley (Washington University, St. Louis, MO) were cultured at 26°C in M199 medium (Sigma-Aldrich, USA), supplemented with 100 units/ml penicillin (Sigma-Aldrich, USA), 100 µg/ml streptomycin (Sigma-Aldrich, USA) and 10% heat-inactivated fetal bovine serum (FBS; Biowest). Whole cell lysate of promastigote was prepared by freeze–thaw cycles as described previously [41].

For axenic amastigotes, the axenically cultured forms grew optimally at a temperature of 32-33°C in a growth media with pH of 5.4. The axenic amastigotes were prepared according to the standard protocol. Briefly, the late-log promastigotes were adapted in an acidic media (RPMI-1640/25 mM 2-(N-morpholino) ethane sulfonic acid (MES)/pH 5.5), at 26°C. These parasites were then grown in RPMI-1640/MES/pH 5.5 at 37°C with 5% CO2.

### Stable transfection of the parasite Leishmania

*Leishmania donovani* Bob strain (*Ld*Bob strain/MHOM/SD/62/1SCL2D) promastigotes were transfected with pXG-GFP-Sphka OE plasmid which harboured the neomycin resistance cassette using electroporation technique as described previously [42] with a Bio-Rad Gene Pulser device and cultured at 26°C in M199 medium (Gibco, Thermofisher scientific USA), supplemented with 100 units/ml penicillin (Sigma-Aldrich, USA), 100 µg/ml streptomycin (Sigma-Aldrich, USA) and 10% heat-inactivated fetal bovine serum (FBS; Gibco, USA). A final concentration of 250 μg/mL of G418 (Sigma-Aldrich, USA) was used to select transfected parasites.

### THP-1 cell culture and infection

THP-1 cells, an acute monocytic leukaemia-derived human cell line (202 TM; American Type Culture Collection, Rockville, MD), were maintained in RPMI-1640 (Sigma-Aldrich, USA) medium supplemented with 10% heat-inactivated FBS (Biowest, UK), 100 units/ml penicillin and 100 μg/ml streptomycin at 37°C and 5% CO_2_. Cells (10^6^ cells/well) were treated with 50 ng/ml phorbol-12-myristate-13-acetate (PMA) (Sigma-Aldrich, USA) for 48h to induce differentiation into macrophage-like-cells before infection. Cells were washed once with phosphate-buffered saline (PBS) and incubated in RPMI medium (Sigma-Aldrich, USA), 10% heat-inactivated FBS, 100 units/ml penicillin and 100 μg/ml streptomycin, before infection. To carry out *in vitro* infection assays, late stationary phase promastigotes (WT) were used at a ratio of 20 parasites per macrophage. *Leishmania*-infected macrophages were incubated at 37°C in a 5% CO_2_-air atmosphere for 4h to allow the establishment of infection and proliferation of intra-macrophage parasites. The cells were then washed five times with PBS to remove non-adherent extracellular parasites. After that, the cells were incubated in RPMI medium at 37°C in a 5% CO_2_-air atmosphere for 48h.

### Infectivity assay

THP-1 cells (1 × 10^6^ cells/well), treated with 50 ng/ml of PMA (Sigma-Aldrich, USA) were seeded on glass coverslips in 6-well plates for 48h. They were infected and simultaneously treated with inhibitors as described above, and the intracellular parasite load (mean number of amastigotes per macrophage) was visualized by Giemsa staining as described previously by Pawar *et al* [43] and expressed as a percentage of the blank controls without drug. At least 10 fields were counted manually for each condition to determine the average number of parasites per macrophage.

### qRT-PCR for gene expression analysis

Total RNA from infected macrophages was isolated using the TRIZOL reagent (Sigma-Aldrich, USA) and its concentration was determined by Nanodrop (Thermo Fischer, USA). cDNA was prepared from one microgram of RNase-free DNase treated total RNA using first-strand cDNA Synthesis Kit (Thermo Fischer Scientific, USA), as per manufacturer’s instructions, using random hexamer primers. The resulting cDNA was analyzed by quantitative real-time (qRT-PCR) RT-PCR (Applied Biosystems, 7500 Fast Real-Time PCR System, CA, USA) with gene-specific primers using PowerUp SYBR Green PCR Master Mix (Thermo Fisher Scientific, USA). The details of the primers (sequences and annealing temperatures) are given in **Table S1.** Thermal profile for the real-time PCR was amplification at 50°C for 2 min followed by 40 cycles at 95°C for 15 sec, 60°C for 1 min and 72°C for 20 sec. Melting curves were generated along with the mean C_T_ values and confirmed the generation of a specific PCR product. Amplification of RNU6AP (RNA, U6 small nuclear 1; THP-1 cells) was used as internal control for normalization. The results were expressed as fold change of control (Uninfected samples (RNU6AP)) using the 2^-ΔΔ*CT*^method. All samples were run in triplicates, including a no-template (negative) control for all primers used.

### Western blotting

Western blot analysis was done as described previously by Darlyuk et al., 2009 [44]. Briefly, protein was isolated from *Leishmania* infected macrophages by resuspending cell lysates in RIPA buffer. Before lysis, adherent cells were placed on ice and washed with PBS. Macrophages were scraped in the presence of RIPA lysis buffer containing 1% NP-40, 50 mM Tris-HCl (pH 7.5), 150 mM NaCl, 1 mM EDTA (pH 8), 10 mM 1,10-phenanthroline and phosphatase and protease inhibitors (Roche). After incubation, lysates were centrifuged for 15 min to remove insoluble matter.The proteins in the lysates were quantified, 80μg of the lysate was boiled (95°C) for 5 min in SDS sample buffer and was subjected to electrophoresis on a 10% SDS-polyacrylamide gel. Proteins were then transferred onto nitrocellulose (NC) membrane using an electrophoretic transfer cell (Bio-Rad Laboratories, USA) at RT. The membrane was washed with 1× TBST solution three times and blocked with 5% BSA for 2h at RT. The blocked membrane was washed in TBST solution three times. The membrane was then incubated with primary monoclonal antibody Caspase 9 (NBP1-69235) in PBS-Tween 20 containing 5% BSA and incubated overnight at 4^ο^C. The blots were subsequently incubated with the secondary antibody conjugated to horseradish peroxidase at 1:3000 dilution in 5% PBS-Tween 20 for 2h at RT. Enhanced chemiluminescence reaction was used for the detection of the blot. The results were expressed as fold change and quantitated by using AlphaEaseFC image analysis software (Alpha Innotech). The data were expressed as mean ± SD of three independent experiments, and the representative image of one experiment is shown.

### Localization of LdSphK1 in L. donovani promastigotes

For detection of SphK1 localization in parasites, 5*10^6^ log phase *Leishmania donovani* promastigotes were plated on poly-L-Lysine coated glass coverslips followed by treatment with 500nM PF-543 diluted in incomplete M199 medium for 6h at 22°C in dark. After 6h of treatment, the cells were washed with 1XPBS, fixed (PBS, 4% formaldehyde, 30 min) and permeabilized (PBS, 0.5% Triton X-100, 5 min). Slides were blocked (1% BSA, 1XPBS, overnight at 4°C or for 30 minutes on rocker, RT). Next day blocked cells were incubated with mice raised anti-SphK1 primary antibody (1:500) for 2 hours at RT. The cells were then washed three times with blocking buffer followed by incubation with Alexa 488 conjugated secondary antibody for 1 hour at RT. The coverslips were mounted on glass slides with antifade DAPI solution for visualization under a confocal laser scanning microscope (Olympus FluoViewTM FV1000 with objective lenses PLAPON ×60 O, NA-1.42) at an excitation wavelength of 556 nm. Images were processed via NIS-Elements software version 4.50. The mean fluorescence intensities were plotted using GraphPad Prism version 8.0. The experiment was performed thrice.

### NBD-Sph assay

A fluorescent molecule: omega (7-nitro-2–1, 3-benzoxadiazol-4-yl [2S,3R,4E]-2-amino octadec-4-ene-1,3-diol [NBD–Sphingosine (NBD-Sph); Avanti Polar Lipids]) was used as a substrate and conversion of NBD-Sph to NBD-S1P was evaluated where SIP was estimated as the end metabolite of the sphingolipid metabolism. Towards this, *rLd*SphK1 protein was incubated with 10μM NBD-Sph, at 37°C for 60 min. After incubation, the recombinant protein was lysed with RIPA lysis buffer to obtain host-free protein. The recombinant protein was thoroughly mixed with 260μL of methanol and 400μL of chloroform:methanol (1:1) followed by adding 16μL of 7 M NH4OH, 400μL of chloroform, and 300μL of 1.5 M KCl. The lipids were separated by centrifugation at 17,000 × g for 5 min. A 100μL aliquot of the upper (aqueous) phase was transferred to a black 96-well flat-bottom plate (Corning).

Fluorescence intensity of the aqueous phase containing NBD-S1P, in *rLd*SphK1 protein sample was measured at an excitation/emission wavelength of 485 nm/530 nm using Varioskan LUX Multimode Microplate Reader (ThermoFisher Scientific). The fluorescence intensity of NBD-S1P was measured.

### Checkerboard assay

Checkerboard assays were put to evaluate the effects of the combination of Amphotericin-B (anti-fungal agent) and PF-543 (Sphingosine kinase inhibitor) against the *Leishmania* parasite. For this, log-stage promastigotes (42–44h) maintained in M199 media were dispensed in a 96-well plate. Amphotericin-B was added vertically at different concentration ranges while PF-543 was added horizontally at different drug concentration ranges in 8*8 format. As a result, the checkerboard consists of columns and rows in which each of the well along the x-axis contains drug Amphotericin-B at different concentrations (12.5nM, 30nM, 50nM and 100nM) and that along the y-axis contains PF-543 at different concentrations (150nM, 250nM, 500nM and 2µM). The plate was incubated at 25°C in a humidified chamber for 48h. The fractional inhibitory concentration (FIC index = FIC A + FIC B, where FIC A is the IC_50_ of drug (A) in combination/IC_50_ of drug A alone, and FIC B is the IC_50_ of drug B in combination/IC_50_ of drug B alone) of each drug was calculated and plotted as an isobologram. A straight diagonal line with an FIC index equal to 1 indicates an additive effect between drug A and drug B, a concave graph below the diagonal with an FIC index of less than 1 indicates a synergistic effect, and a convex curve above the diagonal with an FIC index of more than 0 indicates antagonism [45–47].

### Estimation of reduced (GSH) and oxidized (GSSG) glutathione levels

The estimation of glutathione levels in reduced and oxidized forms was performed in accordance with the previously documented protocol [48]. Briefly, *Leishmania* infected macrophages were incubated with varying concentrations of SphK inhibitors for 24h and lysate was prepared. For GSH estimation, the lysate was added to an assay mixture containing 0.1 mM sodium phosphate buffer (pH 8) and o-phthalaldehyde (1 mg/ml in methanol, Sigma Aldrich, USA) and incubated at RT for 15 min. GSH level was estimated by monitoring fluorescence change at 365Exc/430Emi using a microplate reader. GSSG was also estimated using the same method, however, the lysate was incubated first with N-ethylmaleimide (Sigma Aldrich, USA) in the dark for 5 min, to block the GSH to GSSG oxidation. Results plotted as redox index (GSH/GSSG) ratio are the mean of three independent experiments.

### Animal Studies

Swiss mice were housed under standard conditions of food, temperature (25 ± 3 °C), relative humidity (55 ± 10%) and illumination (12h light/dark cycles) obtained from the Central Laboratory Animal Resources, Jawaharlal Nehru University, Delhi.

For experimental assays, infected mice were divided into 4 groups, each group comprise of 3 mice. The uninfected group of mice was used as an experimental negative control. The experimental group of mice were infected with *L. donovani* 24h prior to injection with 10 mg/kg PF-543 and 2mg/kg amphotericin B alone via I.P route for 3 consecutive days at a dose of body weight, as a Control. While evaluating the synergistic formulation, comprising 2mg/kg PF-543 and 0.4mg/kg amphotericin B, another set of experimental mice were treated once per day for three consecutive days. Evaluation of the immunomodulatory potential was done using cytokine profiling in *Leishmania* infected Swiss mice treated with PF-543 and Amphotericin-B alone and simultaneously followed by determination of parasite infectivity using *in-vitro* parameters. To evaluate the parasitemia and expression of inflammatory markers in infected mice upon drug administration, spleen was harvested from each group of mice and qRT-PCR analysis was performed using primers specific for inflammatory markers; TNF-α and IL-10 and parasite specific kinetoplast minicircle gene; JW in PF-543 and Amphotericin-B treated infected mice in respect to the untreated Swiss mice.

### Ethics statement

Animal studies were performed following CPCSEA guidelines and approved by the Institutional Animal Ethics Committee (IEAC) of JNU. Swiss mice were obtained from the Central Laboratory Animal Resources, JNU, New Delhi, and maintained under standard conditions.

### Statistical analysis

Fold-expression (qRT-PCR and densitometric analysis), NBD-Sph and intracellular parasite burden were represented as mean ±SD. Each experiment was repeated three times in separate sets. Statistical differences were determined using Student’s unpaired 2-tailed *t*-test. All statistics were performed using GraphPad Prism Version 5.0 (GraphPad Software, USA). p ≤ 0.05 was considered significant [* (P<0.01 to 0.05), ** (P< 0.001), *** (P< 0.0001), ns (P≥ 0.05)].

## Results

### 1. Expression and Purification of the recombinant *Leishmania* SphK1 (*rLdSphK1*) protein and its Functional Characterization in presence and absence of SphK1 inhibitor-PF-543

Detection of *Leishmania-*specific Sphk-1 (*Ld*SphK-1) activity in promastigotes was evaluated using generic inhibitors of SphK; 500nM PF-543, 500μM DMS and 100μM SKI-5C (IC_50_; **Fig 4, Fig S3 (A,B)**) for 48h. To investigate the impact of SphK inhibitors; such as PF-543, DMS and SKI-5C on catalytic activity of SphK1 expressed by *Leishmania* promastigotes (*Ld*SphK1), we performed fluorometric assay based estimation of enzymatic activity using 7-nitro-2-1,3-benzoxadiazol-4-yl (NBD)-tagged sphingosine (NBD-sphingosine). The results demonstrated significant reduction of NBD-S1P levels in PF-543-treated *Leishmania* promastigotes, in comparison to untreated control **(Fig 1A),** validating the active catalytic form of the *Ld*SphK1.

**Fig 1.**
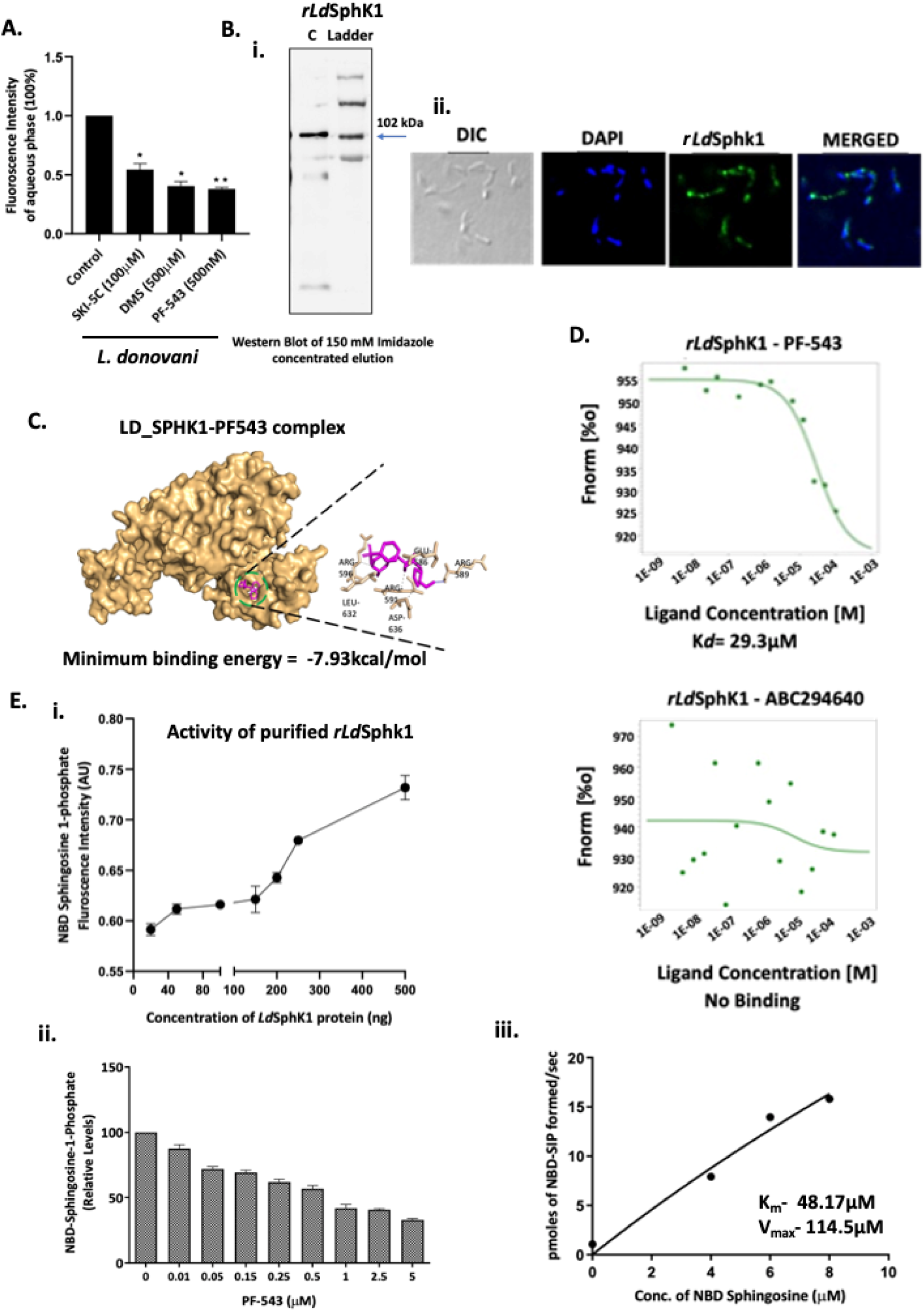
(A) Estimation of *Ld*SphK-1-mediated generation of NBD-SIP levels against SphK inhibitors in *L. donovani* promastigotes. *L. donovani* bob promastigotes were cultured in 25 cm^2^ flasks followed by treatment with SphK inhibitors; PF-543, DMS and SKI-5C for 48h. Inhibition of SphK1 using these SphK inhibitors cause decreased S1P levels. SphK inhibitor treated promastigotes were resuspended in buffer containing fatty acid-free BSA (0.1% (w/v)) followed by resuspension in buffer containing BSA (1% (w/v)) and incubated with NBD-sphingosine (10μM) for 45 min at 37 °C. Promastigotes incorporate NBD-sphingosine (NBD-Sph), which are phosphorylated by Sphingosine kinase (SphK1) to NBD-S1P. Bar graph depicts ELISA-based S1P quantification in promastigotes treated with inhibitors. **(B) Purification and sub-cellular localization of Recombinant *rLd*SphK1 in *L. donovani***. Concentrated r*Ld*SphK1 protein. (i) Gel image showing a purified band of 102kDa recombinant *Ld*SphK1 (ii) The subcellular localization of recombinant *Ld*SphK1 was investigated in promastigotes. Panel A: DIC at 60X. Panel B: DAPI. Panel C: anti-*Ld*SphK1 antibody detected using FITC (green)-conjugated secondary antibody. Panel D: merged micrographs. *Ld*SphK1 expression represented in the green channel. Co-localization shown as the merged image. **(C) *In silico* docking of the *rLdSphK1* in presence of PF-543.** *In-silico* ligand-substrate interaction was done using Autodock 1.5.7rc1 and Cygwin terminal. The *in silico* docking generated the complex of *rLdSphK1*-PF-543 and represented as surface model highlighted with residues involved in the interaction using Chimera, Ligplus, Discovery Studio v19.1.0, and PyMOL v2.3.2 software. **(D) Biophysical interaction and functional characterization of the *rLdSphK1* (20µM) in presence of PF-543 (500nM, 1µM, 2.5µM, 5µM, 10µM, 20µM).** MST analysis confirms the interaction between *rLdSphK1* and PF-543. Dose-response curves of *rLdSphK1*–PF-543 showed K_D_ of 29.3µM. **(E) Analysis of functional characterization and catalytic activity of the *Ld*SphK1 recombinant protein.** (i) Time kinetics of activity of purified *rLd*SphK1 and NBD-S1P-based fluorometric assay showing increase in NBD-SIP levels in presence of different concentrations of *rLd*SphK1 protein. (ii) The inset shows inhibition of NBD-SIP levels in the presence of different concentrations of PF-543 using 200ng of recombinant *Ld*SphK1. (iii) Varying different concentration of NBD Sphingosine (2-10μM) and keeping *rLd*SphK1 constant (200ng), NBD-SIP levels were measured to calculate K_m_ and V_max_ of *rLd*SphK1. The plate was immediately placed on the varioskan LUX Multimode Microplate Reader (Thermo fisher, Massachusetts, USA) every 5 seconds for 20 min.

Next, *Ld*Sphk1 was successfully cloned in pET-28 a(+) vector with N-terminal 6x-Histidine-tag **(Fig S1B)** and recombinant protein was purified using affinity based purification method **(Fig S1C)**. For this, we went for expression and purification of the recombinant *Ld*SphK1 (*rLd*SphK1) protein and observed a single band corresponding to ∼102 kDa on SDS PAGE **(Fig 1B(i))** corresponding to *Leishmania* specific SphK1. Further, the subcellular localization of *rLd*SphK1 was investigated in *L. donovani* Bob cells using confocal microscopy for validation of in-house generated *rLd*SphK1 antibody. Immunostaining of *L. donovani* using anti-*Ld*SphK1 antibody suggested cytoplasmic or membranous localization of *rLd*SphK1 **(Fig 1B(ii)).**

Furthermore, we conducted computational docking analyses of the *Ld*SphK1 protein with the PF-543. The optimal conformations of the docked compounds were chosen based on their minimum free binding energy to the binding domain. Upon analysis of this pocket, it was discovered that there is a single hydrogen bond between residue ARG589 and ligand, with a distance of 2.03 Å. In addition, GLU586, ARG591, ARG596, LEU632, and ASP636 established hydrophobic contacts within the pocket. The binding of PF-543 to the protein is facilitated by these interactions, which result in a minimum binding energy of -7.93 kcal/mol **(Fig 1C)**. The analysis of these results indicates that PF-543 creates a stable structure when bound to the *Ld*SphK1 binding pocket. These data inferred that *rLdSphK1* and PF-543 are possible interacting partners having favourable binding energy.

To validate the biophysical interaction between purified *rLd*SphK1 and PF-543 and analyze the binding and kinetics, Microscale Thermophoresis (MST) was performed in NanoTemper Monolith NT.115 instrument. Herein, labelled *rLdSphK1* (20µM) was titrated against varying concentrations of PF-543 (500nM, 1µM, 2.5µM, 5µM, 10µM, 20µM). The dose response analysis for *rLdSphK1*-PF-543 binding revealed dissociation constant, K_D_ to be 29.3µM, suggesting strong binding of *rLdSphK1* with PF-543 **(Fig 1D).** Additionally, we also evaluated interaction between *rLdSphK1* and ABC294640 as a negative experimental control where no such binding between the two could be detected.

The time course kinetics for *rLdSphK1* was done using NBD-Sphingosine based fluorometric assay for detection of NBD-S1P levels. The results represented a steep increment in NBD-S1P level with *rLdSphK1* concentration ranging from 200ng to 500ng **(Fig 1E(i)),** following which saturation in activity was detected (data not shown). In addition, the inhibition of *rLd*SphK1 catalytic activity was also evaluated in a dose dependent manner with PF-543’s concentration ranging from 1 to 5µM. The results represented a maximum inhibition at 5µM of PF-543, while the protein concentration was kept constant at 200ng **(Fig 1E(ii)).** To detect Km and Vmax values, varying concentration of NBD Sphingosine (2-10µM) was taken along with 200 ng of *rLd*SphK1. The results confirmed 48.17μM and 114.5μM as Km and Vmax respectively, according to Lineweaver-Burk plots, while D-erythro-sphingosine is used as the substrate for *Ld*SphK1 **(Fig 1E(iii)).**

### 2. Expression and Purification of the recombinant Human SphK1 (*rhSphK1*) protein and its Functional Characterization in presence and absence of SphK1 inhibitor-PF-543

Initially, we conducted the amino acid sequence alignment of LD_SphK1 and Human_SphK1 utilizing MEGAX program (version 10.2.2). It demonstrated considerable alignment with similar and a limited number of identical amino acids **(Fig S4A)**. Additionally, we conducted 3D structural alignment of the two protein structures with Pymol software. We observed substantial alignment between the two proteins, with an RMSD of 0.585. Furthermore, the alpha helix and beta sheets of the Human DAGKc domain (Blue) are accurately aligned with the LD_DAGKc domain (Yellow) in **(Fig S4B)**. A phylogenetic tree was created utilizing SPHK1 gene sequences from various species of *Leishmania*, humans, rats, and mice. The phylogenetic tree was constructed using neighbor-joining technique with MEGAX software. Bootstrap analyses with 1000 replicates were conducted to evaluate the robustness of the constructed phylogenetic tree **(Fig S4C)**.

Next, *Human*Sphk1 was successfully cloned in pET-28 a(+) expression vector, and the recombinant plasmid was transformed into *E. coli* strain BL21 (LDE3) gold. Later, we went for expression and purification of the recombinant *Human*SphK1 (*rHuman*SphK1) protein **(Fig S2B).** A single band corresponding to ∼50 kDa was observed on SDS PAGE **(Fig 2A)** corresponding to Human specific SphK1.

**Fig 2.**
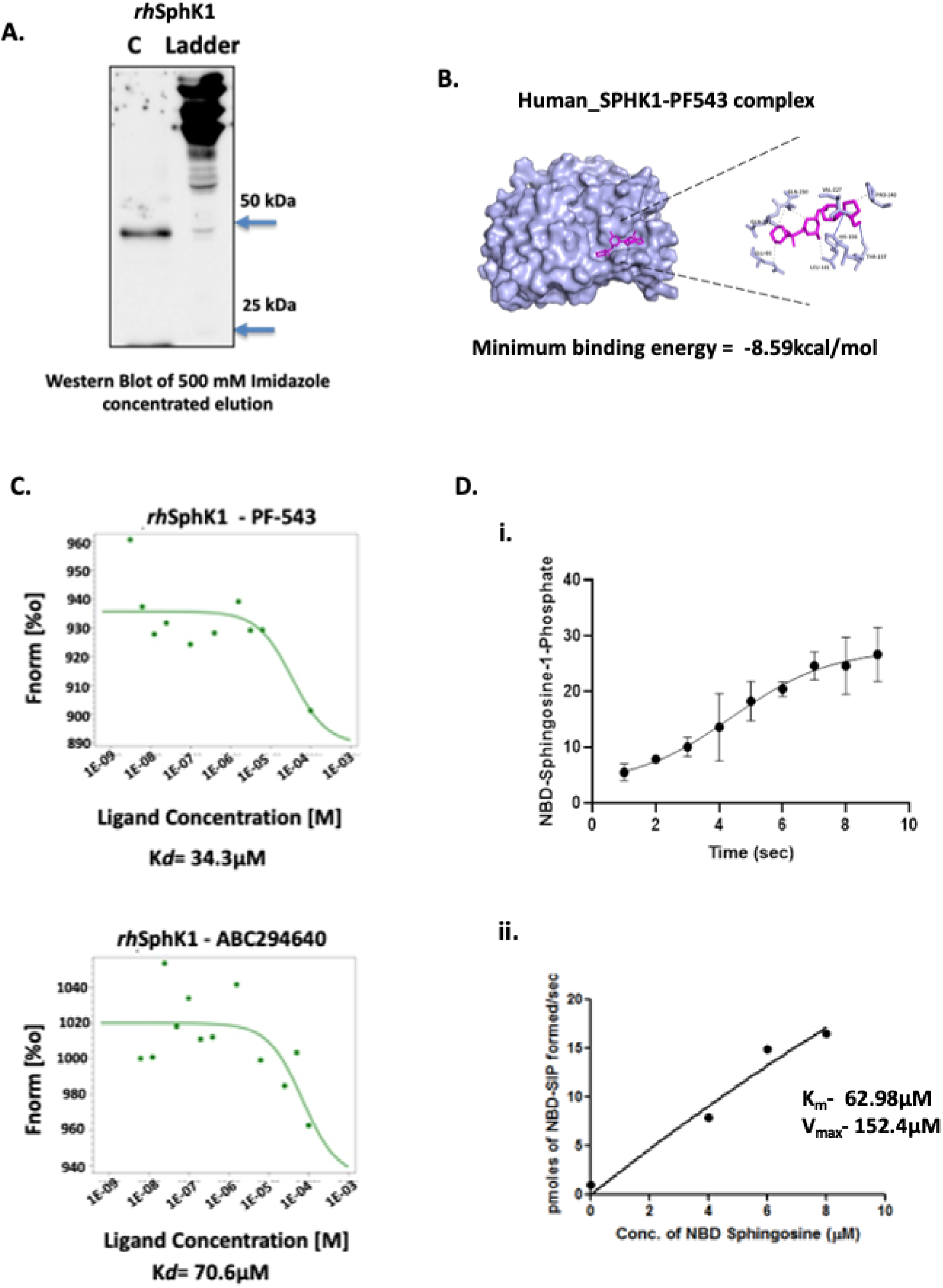
(A) Purification of Recombinant *rhSphK1* from Supernatant. Gel image showing a concentrated *rhSphK1* protein band of 50kDa. **(B) *In silico* docking of the *rHumanSphK1* in presence of PF-543.** *In silico* ligand-substrate interaction was done using Autodock 1.5.7rc1 and Cygwin terminal. The *in silico* bound complex of *rhSphK1*-PF-543 was represented as surface model highlighted with residues involved in the interaction using Chimera, Ligplus, Discovery Studio v19.1.0, and PyMOL v2.3.2 software. The list of residues involved in hydrophobic interaction and hydrogen bond formation were highlighted in the model. **(C) Functional characterization of the *rhSphK1* in presence of PF-543.** MST analysis confirms the biophysical interaction between *rhSphK1 **-***PF-543 and *rhSphK1-* ABC294640, which was used as a control. Dose-response curves of PF-543 (20µM) and ABC294640 showed K_D_ of 34.3µM and 70.6µM respectively. **(D) Analysis of catalytic activity of the *rhSphK1* using NBD-S1P-based fluorometric assay. (i)** Time kinetics of activity of purified *rhSphK1* demonstrated increase in NBD-SIP levels in presence of 200ng of *rhSphK1* protein. **(ii)** Varying different concentration of NBD Sphingosine (2-10μM) and keeping *rhSphK1* constant (200ng), NBD-SIP levels were measured to calculate K_m_ and V_max_ of *rhSphK1*. The readings were recorded every 5 seconds for 20 min using varioskan LUX Multimode Microplate Reader (Thermo fisher, Massachusetts, USA).

In order to assess the underlying physiological mechanisms of protein-ligand interactions, we conducted *in-silico* docking experiments involving the hSphK1 protein and the PF-543 compound. The optimal conformations of the docked compounds were chosen based on their minimum free binding energy to the binding domain. Upon examination of this pocket, it was discovered that it contains two residues, namely HIS156 and THR157, which are linked together by hydrogen bonds. The hydrogen link between them has a distance of 3.44 Å, while the other hydrogen bond has a distance of 2.53 Å. Similarly, GLU93, LEU161, VAL227, GLN230, GLN231, and PRO240 established hydrophobic interactions within the pocket. The binding of PF-543 to the protein is facilitated by these interactions, resulting in a minimum binding energy of -8.59 kcal/mol **(Fig 2B).** The analysis of these results indicates that PF-543 creates a stable structure when bound to the hSphK1 binding pocket. These data inferred that *rhSphK1* and PF-543 are possible interacting partners having favourable binding energy.

To validate the biophysical interaction between purified *rhSphK1* and PF-543, MST was performed in NanoTemper Monolith NT.115 instrument. Herein, labeled *rhSphK1* (20µM) was titrated against varying concentrations of PF-543. The dose response analysis for binding between *rhSphK1*-PF-543 and *rhSphK1*-ABC294640 (SphK2 inhibitor) revealed dissociation constant, K_D_ to be 34.3µM, and 70.6µM respectively suggesting strong binding of *rhSphK1* with PF-543 and ABC294640 respectively **(Fig 2C).**

Further, the time course kinetics for *rhSphK1* was done using NBD-Sphingosine based fluorometric assay for evaluating its functional active form. The results represented a steep increment in generation of NBD-S1P level with 200 ng *rhSphK1* **(Fig 2D (i)),** following which saturation in activity was detected (data not shown). To detect Km and Vmax values, varying concentration of NBD Sphingosine (2-10µM) was taken along with 200ng of *rhSphK1*. The results confirmed 62.98μM and 152.4μM as Km and Vmax respectively with D-erythro-sphingosine is used as the substrate for *rhSphK1* **(Fig 2D (ii)).**

### 3. Effect of PF-543 on cytotoxicity and infectivity of *L. donovani* SphK1 overexpressor *(Ld*SphKa) promastigotes

To confirm successful transfection of *Leishmania donovani* promastigotes with the pXG-GFP-SphKa construct, parasites were examined under confocal microscopy for GFP expression. A distinct population of GFP-positive parasites was detected, confirming transgene expression, alongside a subset of GFP-negative cells **(Fig 3A).** Non-transfected GFP-tagged parasites were included as controls to validate the specificity of transgene expression.

**Fig 3.**
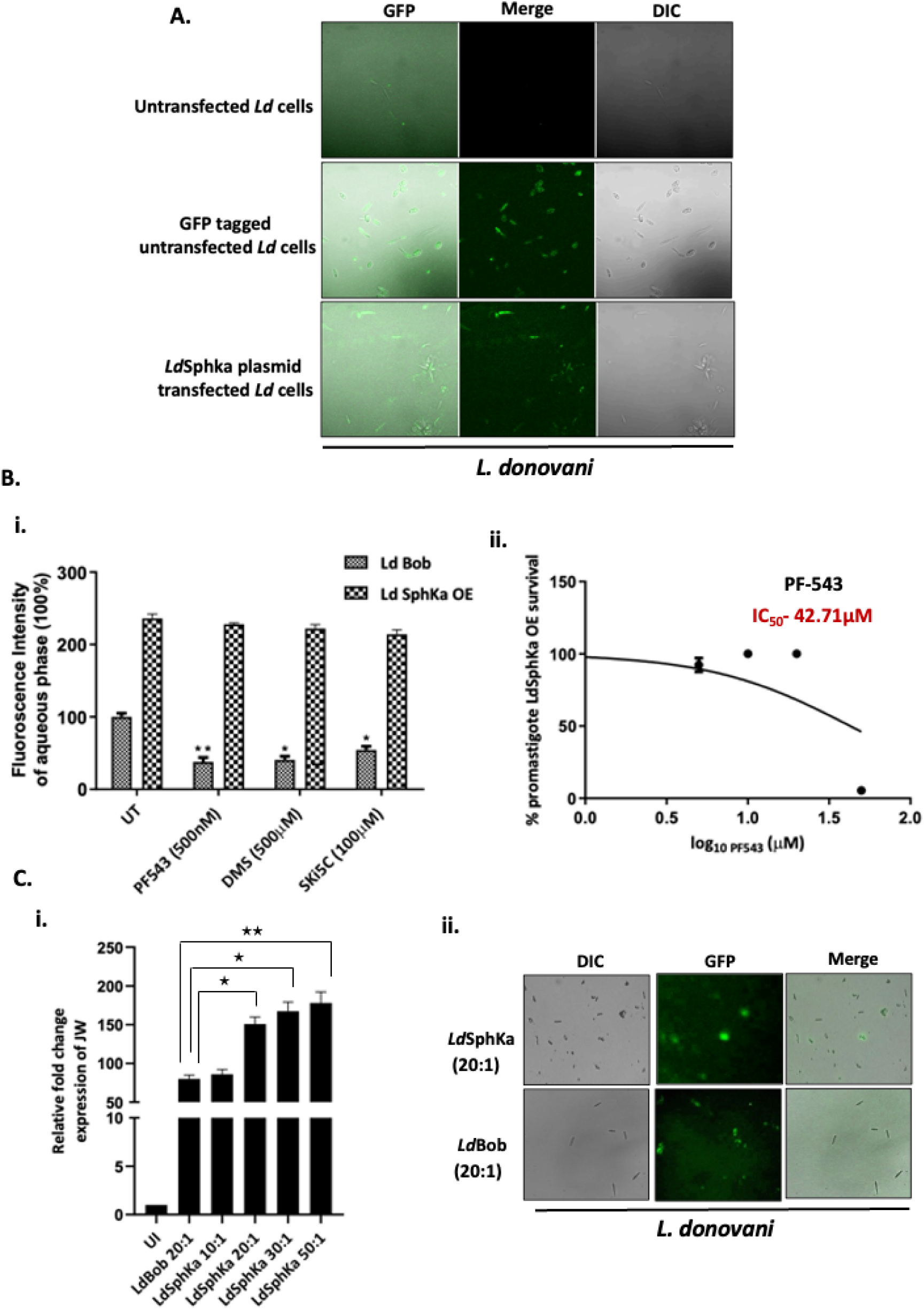
Effect of PF-543 on cytotoxicity and infectivity of *L. donovani* SphK1 overexpressor promastigote cell line. **(A)** Detection of the presence of GFP encoded SphKa plasmid in *Leishmania donovani* parasites. *L. donovani* bob promastigotes were cultured in 25 cm^2^ flasks followed by transfection with GFP-tagged SphKa plasmid. **(B (i)) Estimation of *Ld*SphK-1-mediated generation of NBD-SIP levels against PF-543 in *L. donovani* SphKa promastigotes.** *L. donovani* bob promastigotes were cultured in 25 cm^2^ flasks followed by treatment with SphK1 inhibitor; PF-543 for 48h. Inhibition of SphK1 using PF-543 had no significant effect on S1P levels. SphK inhibitor treated SphKa promastigotes were resuspended in buffer containing fatty acid-free BSA (0.1% (w/v)) followed by resuspension in buffer containing BSA (1% (w/v)) and incubated with NBD-sphingosine (10μM) for 45 min at 37 °C. Promastigotes incorporate NBD-sphingosine (NBD-Sph), which are phosphorylated by Sphingosine kinase (SphK1) to NBD-S1P. Bar graph depicts ELISA-based S1P quantification in SphKa promastigotes treated with PF-543. **(ii) Effect of SphK1 inhibitor; PF-543 in *Leishmania* overexpressor *spp*.** To evaluate the IC_50_ for PF543, approximately 5×10^4^ *Ld SphKa* Bob cells were seeded in each well of 96-well flat bottom plates and supplemented with M199 media containing PF-543 (200µl/well). The IC_50_ was found to be 42.71μM for *Leishmania donovani* SphKa promastigotes. Each experiment was done in triplicates and repeated thrice. **(C (i)) Effect of infectivity of *L. donovani* SphK1 overexpressor on the clearance of intracellular parasite load.** THP-1 macrophages were cultured in six-well plates in the presence or absence of *L. donovani* infection (MOI, 20:1) for 6h. Further, THP-1 cells were infected with *L. donovani* SphK1 overexpressor promastigotes at various MOIs (10:1, 20:1, 30:1, 50:1). Infected THP-1 were washed to remove non-internalized parasites. Total RNA was enriched using TRIzol and the resulting cDNA was subjected to real-time PCR analysis using primers specific for infectivity (JW) genes in infected macrophages. RNU6A was used as a housekeeping gene. The results are expressed as fold-change of uninfected THP-1 cells. Statistical significance was quantified using the unpaired t-test. The data is a representation of mean ± SD from three independent experiments *, p< 0.05; ***, p < 0.001. **(C (ii)) Analysis of GFP expression in *L. donovani* SphK1 overexpressor and wild type promastigote cell lines.** Detection of the presence of GFP encoded SphKa and *Bob* plasmids in *Leishmania donovani* and *Leishmania* overexpressor parasites. *L. donovani bob* and *L. donovani* overexpressor promastigotes were cultured in 25 cm^2^ flasks followed by confocal microscopy.

Detection of *Leishmania*-specific Sphk-1 (*Ld*SphK-1) activity in *Leishmania* SphK1 over-expressor promastigotes was detected using generic inhibitors of SphK. To investigate the impact of SphK inhibitors; such as PF-543, DMS and SKI-5C on catalytic activity of SphK1 expressed by *Leishmania* over-expressor promastigotes (*Ld*SphKa), we performed fluorometric assay based estimation of enzymatic activity using 7-nitro-2-1,3-benzoxadiazol-4-yl (NBD)-tagged sphingosine (NBD-sphingosine). The results demonstrated higher but similar levels of NBD-S1P in PF-543, DMS and SKI5C-treated *Ld*SphKa promastigotes, in comparison to untreated control **(Fig 3B (i)),** validating the overexpression of active catalytic form of the SphK1 in *Leishmania donovani*.

Subsequently, the cytotoxic effect of PF-543 on *Leishmania* SphK1-overexpressing promastigotes was assessed using the MTT assay. PF-543 exhibited cytotoxicity against *Ld*SphKa promastigotes with an ICLL value of 42.71µM, notably higher than that observed for wild-type *L. donovani* promastigotes **(Fig 3B (ii)).** These results suggest that overexpression of SphK1 may confer partial resistance to PF-543-induced cytotoxicity, potentially by modulating downstream lipid signaling pathways.

To evaluate the impact of SphK1 over-expression on clearance of intracellular load of amastigotes, THP-1 cells were infected with *Ld*SphKa promastigotes at various MOIs (10:1, 20:1, 30:1, 50:1). The intracellular parasite burden was analysed using RT-PCR at 48h post-infection. Uninfected (UI)THP-1 cells were used as experimental controls. At an MOI of 10:1, approximately 86% infection was observed in *Ld*SphKa-infected THP-1 cells, indicating efficient parasite entry and survival. In comparison, *Bob*-infected THP-1 cells demonstrated ∼80% infection at an MOI of 20:1 **(Fig 3C).** Across higher MOIs (20:1, 30:1, and 50:1), infection rates remained consistently higher in the *Ld*SphKa group relative to the *Bob*-infected controls **(Fig 3C),** confirming that SphK1 overexpression is associated with enhanced infectivity and a rapid-growth phenotype.

These findings collectively suggest that SphK1 plays a crucial role in promoting parasite survival and proliferation, and that its overexpression may attenuate the antiparasitic effects of PF-543.

### 4. Cytotoxicity and infectivity of *L. donovani* upon PF-543 treatment

The cytotoxic effect of PF-543 was studied in both THP-1 macrophages; an acute monocytic leukaemia-derived human cell line (202 TM; American Type Culture Collection, Rockville, MD) and *Leishmania* spp (*Ld*Bob strain/MHOM/SD/62/1SCL2D) promastigotes using MTT assay. The selectivity index (SI) representing the ratio of cytotoxicity in THP-1/promastigotes (CC_50_/IC_50_) was determined for PF-543 treatment. The results demonstrated PF-543 to be more toxic to the intracellular parasites than to the host cells. The IC_50_ of PF-543 was 90µM for THP-1 macrophages, 17.54µM for intracellular amastigotes and 45µM for axenic amastigotes respectively, as determined by cytotoxic or metabolic viability analysis. These findings are summarized in **Fig 4** that provided information on the Inhibitory Concentrations (IC_50_). The data analysis showed PF-543 had an IC_50_ of 500nM and SI of 180nM **(Fig 4A).** In contrast, ABC294640 was found to be toxic to host cells expressing the SphK2 isoform, with ICLL values of 60µM for THP-1 macrophages and 69.7µM for intracellular amastigotes **(Figure S3C).**

To evaluate the impact of PF-543 on clearance of intracellular load of amastigotes, THP-1 cells were treated with PF-543 (90µM) prior to infection with *L. donovani* at an MOI of 20:1. The intracellular parasite burden was analyzed using RT-PCR and Giemsa staining at 48h post-infection. On analysis, Untreated-uninfected (UT-*Ld*) THP-1 cells were used as experimental controls and could able to show ∼ 80% infection with *L. donovani*. While PF-543 treated THP-1 (24h post-treatment) showed ∼ 60% of infection **(Fig 4B (i,ii)).** Upon comparing the clearance of parasitic load in PF-543 treated infected THP-1 cells (PF543+*Ld*) with untreated infected THP-1 (UN+*Ld*), ∼40% reduction in the number of parasites (amastigotes/macrophage) was detected per infected macrophage **(Fig 4B (i,ii)).**

**Fig 4.**
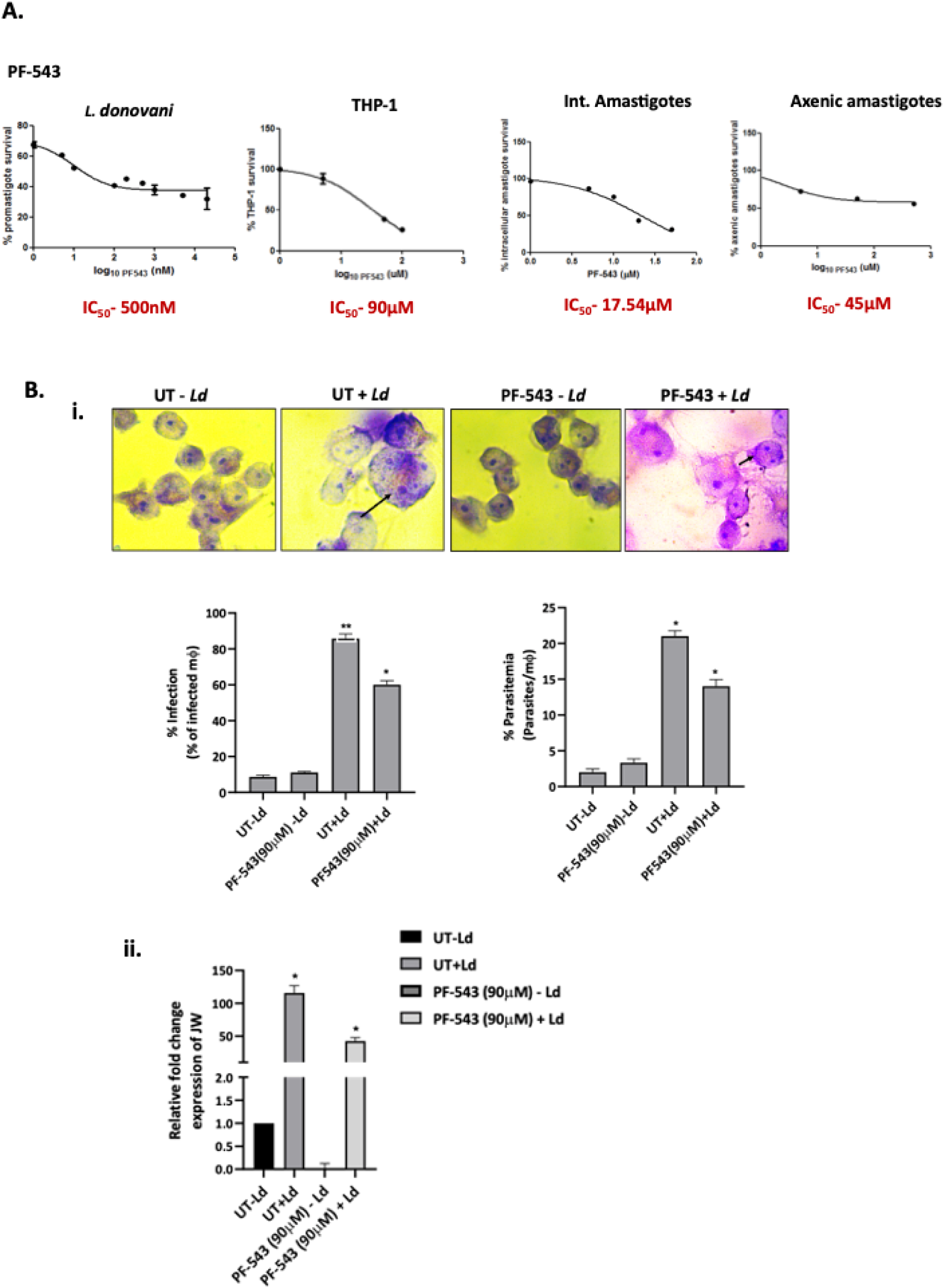
(A) Effect of SphK1 inhibitors; PF-543 and ABC294640 in *Leishmania spp* and host macrophages. To evaluate the IC_50_ for PF543, approximately 6×10^3^ THP-1 and 5×10^4^ *Ld* Bob cells were seeded in each well of 96-well flat bottom plates and supplemented with RPMI and M199 media containing PF-543 (200µl/well) respectively. For intracellular amastigotes, 1 × 10^6^ THP-1 cells, treated with 50 ng/ml of phorbol 12-myristate 13-acetate (PMA) were seeded on glass coverslip in a 6-well plate for 48h. They were infected with late log-phase *L. donovani* promastigotes and simultaneously treated with PF-543. The cells were further incubated for 2 days at 37°C and 5% CO_2_. To determine, the intracellular parasite burdens (mean number of amastigotes per macrophage) were microscopically assessed using Giemsa staining. For axenic amastigotes, the axenically cultured forms grew optimally at a temperature of 32-33°C in a growth media with pH of 5.4. The IC_50_ were found to be 500nM, 90μM, 17.54μM and 45μM for *Leishmania donovani* promastigotes, THP-1 macrophages, intracellular amastigotes and axenic amastigotes respectively. Each experiment was done in triplicates and repeated thrice. **(B) Effect of Sphk1 inhibition on parasite infectivity in THP-1 macrophages.** THP-1 cells grown in RPMI medium were treated with SphK1 inhibitor; PF-543 for 24h followed by infection with *L. donovani* promastigotes at an MOI of 20:1 for 48h. (i) After 48h of infection, Giemsa staining was performed to assess the parasite burden. THP-1 cells were fixed, Giemsa stained and amastigotes were counted visually. Virulence capacity was determined by calculating infectivity (left panel) and parasitemia (right panel). Untreated infected THP-1 cells were used as control. (ii) Total RNA was enriched using TRIzol. Infectivity was validated using qRT-PCR with primer specific marker JW as a molecular indicator. Expression of JW mRNA in infected cells inferred parasite load and represented as percentage infectivity. Data analysis was performed using the 2^-ΔΔ^*^CT^*method. The results are representative of three independent experiments. Statisticalsignificance was quantified using the unpaired t-test. The results signify mean ± S.D with n = 3, *P < 0.05.

These findings suggest that inhibition of host SphK1 via PF-543 significantly reduces intracellular parasite load, likely by impairing the parasite’s ability to sustain its intracellular growth. This points to a persistent, slow-growth phenotype of parasites exposed to SphK1 inhibition.

### 5. Decrease in *Leishmania* glutathione levels mediates ceramide-induced parasite death

*Leishmania* infection leads to increase in ceramide generation that further depletes cholesterol from the membrane and disrupts lipid rafts resulting in weak CD40 mediated signalling that leads to increased ERK1/2 phosphorylation and impaired antigen presentation to the T cells, worsening the diseased condition [49].

DL-threo-PPMP, a ceramide analog is known to inhibit the activity of glucosylceramide synthase. Here in this study, it was used to investigate the role of ceramide in *Leishmania* parasite death. Exogenously added ceramide synthase inhibitor, DL-threo-PPMP at concentrations of 20μM exhibited a dose-dependent cytotoxic effect (expressed as growth inhibition) on THP-1 macrophages **(Fig S5).**

THP-1 cells were infected with *L. donovani* at an MOI of 20:1 followed by treatment with DL-threo-PPMP. The glutathione and apoptotic levels were analysed using GSSH/GSG assay and Western blot at 48h post-treatment. On analysis, we found that addition of ceramide synthase inhibitor induced a decrease in GSH/GSSG ratio **(Fig 5A)** and an increase in Caspase 9 levels **(Fig 5B)** in *Leishmania* infected macrophages.

**Fig 5.**
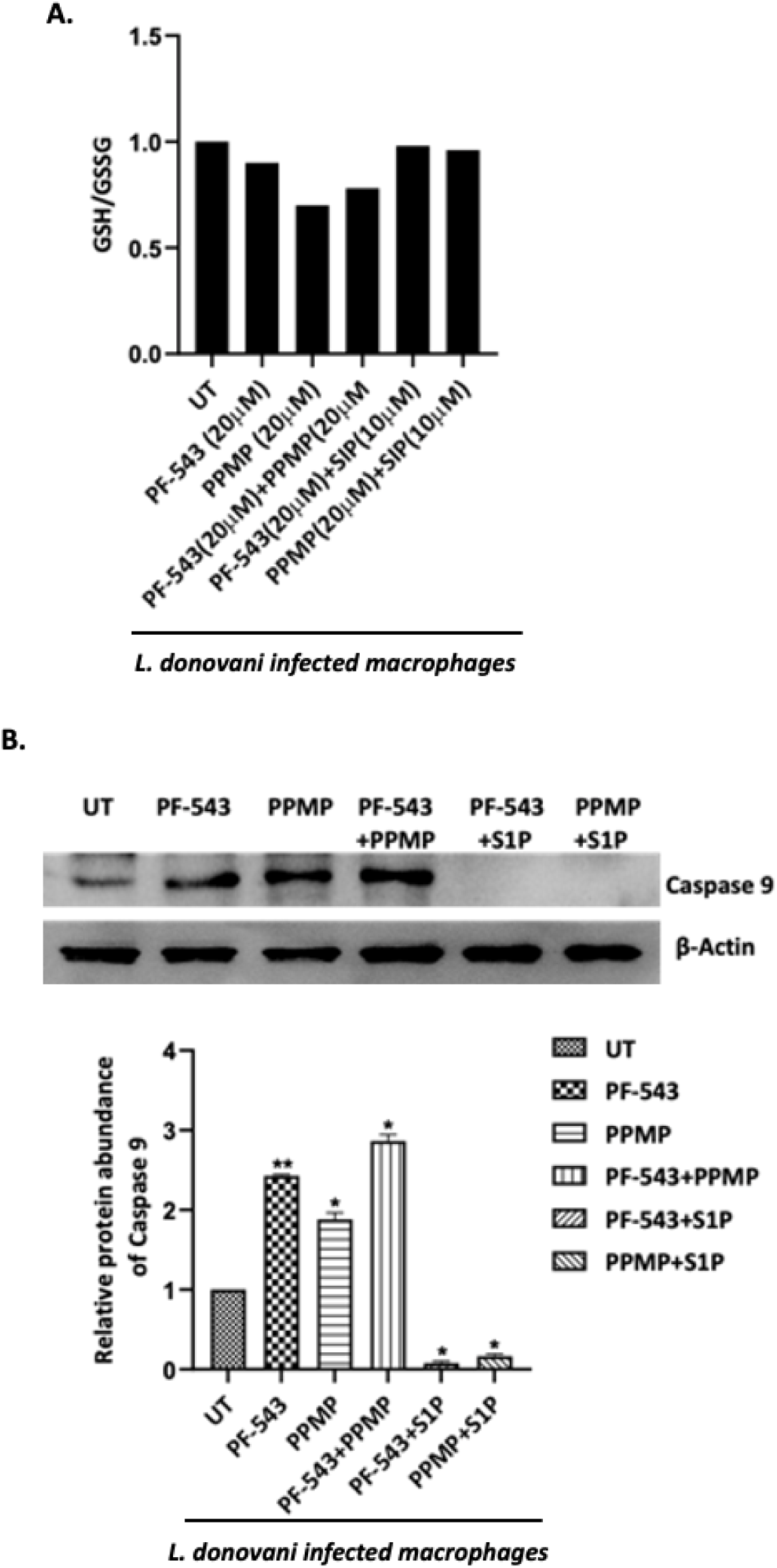
(A, B) Estimation of glutathione and Caspase 9 levels upon PF-543 treatment in *Leishmania* infected macrophages. THP-1 macrophages were cultured in six-well plates in the presence of *L. donovani* infection (MOI, 20:1) for 6h. Infected THP-1 were washed to remove non-internalized parasites and treated with PF-543 (20μM) or DL-threo-PPMP; glucosylceramide synthase inhibitor (20μM) or SIP (10μM) for next 48h. GSH/GSSG and Caspase-9 levels were analysed using cell lysates quantified for GSG/GSSH assay and Western blot. Band intensities were quantified by ImageJ software and were plotted in GraphPad Prism. Data from one of three experiments is shown.

Sphingosine-1-phosphate (S1P), a metabolite of the sphingomyelin pathway and a biological antagonist of ceramide action, prevents ceramide-induced death in mammalian cells [30]. To determine whether S1P can prevent the killing effect of ceramide, *Leishmania* infected macrophages were exposed to PPMP with or without S1P at 10μM (a non-toxic concentration for the parasites). The cytotoxic effect of ceramide found by observing high Caspase levels was significantly reduced by exogenous addition of S1P **(Fig 5B)** and was also found to restore the levels of GSH/GSSG in SIP supplemented cells **(Fig 5A).** These findings suggest the prevention of ceramide-induced toxicity for leishmania parasites by sphingosine-1-phosphate.

### 6. Elucidation of inflammatory responses and infectivity post-PF-543 treatment in *Leishmania* infected macrophages

Cytokines play a crucial role in mediating T-cell responses and host defense mechanisms in macrophages. LPS potently activates macrophages and cytokine signalling by TLR4 stimulation, while *Leishmania* parasites suppress LPS-induced host inflammatory cytokine responses [50]. We investigated the effect of perturbation of the host SphK1 pathway on the pro-inflammatory and anti-inflammatory cytokines from LPS-stimulated *Leishmania* infected macrophages. We found that PF-543 post-treatment in infected macrophages reduced the levels of anti-inflammatory cytokine *IL-10* (^∼^80%) as confirmed by RT-PCR and ELISA while the level of pro-inflammatory cytokines *IL32*γ, *IL-12* and *TNF-*α remain unaltered **(Fig 6A (i,ii)).** We further evaluated the parasite load in presence of PF-543 using qRT-PCR based on estimation of JW (kDNA) mRNA levels in PF-543 treated and untreated infected cells. The results suggested PF-543 post-treatment leads to decrease in parasite load in infected macrophages **(Fig 6B),** thereby indicating the role of PF-543 in reducing the parasite survival.

**Fig 6.**
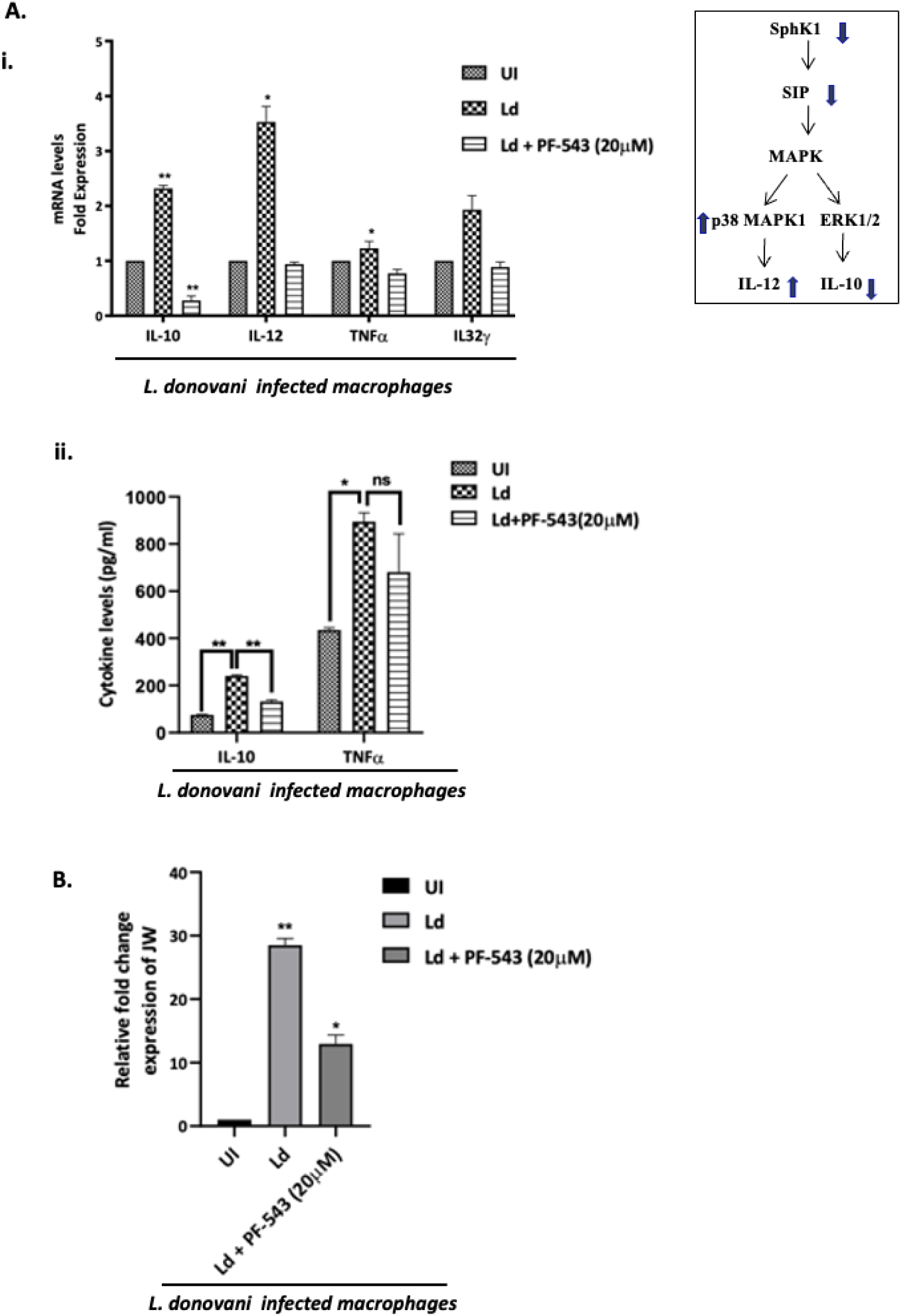
(A(i), B) Effect of PF-543 on the expression of inflammatory molecules and clearance of intracellular parasite load. THP-1 macrophages were cultured in six-well plates in the presence or absence of *L. donovani* infection (MOI, 20:1) for 6h. Infected THP-1 were washed to remove non-internalized parasites and treated with PF-543 for next 48h. Total RNA was enriched using TRIzol and the resulting cDNA was subjected to real-time PCR analysis using primers specific for inflammatory (IL32γ, IL-12, IL-10 and TNF-α) and infectivity (JW) genes in PF-543 treated and untreated infected macrophages. RNU6A was used as a housekeeping gene. The results are expressed as fold-change of uninfected THP-1 cells. Statistical significance was quantified using the unpaired t-test. The data is a representation of mean ± SD from three independent experiments *, p< 0.05; ***, p < 0.001. **(A (ii))** Graph represents cytokine levels measured by sandwich ELISA. Statistical significance was quantified using the unpaired t-test with Welch’s correction, in LPS-stimulated, *Ld*-infected and PF-543 treated macrophages.

### 7. Evaluation of antileishmanial activity of PF-543 and Amphotericin B combined formulation against *L. donovani* promastigotes and intracellular amastigotes

Amphotericin B (Amph-B) is a first-line medication for treating leishmaniasis specifically *Leishmania donovani* infection, in India [51]. Amph-B which binds to the ergosterol of protozoan cells causes a change in membrane integrity resulting in ions leakage, and ultimately leading to cell death [52]. We also checked the potential impact of Amphotericin B the current drug used in clinics, alone and in combination with PF-543 on anti-leishmanial activity against *L. donovani* promastigotes using MTT assay. The *in-vitro* synergistic activity of PF-543 along with Amphotericin B was represented by an isobologram. Upon analysis of antileishmanial activity, Amphotericin B showed an IC_50_ of 30nM and PF-543 showed IC_50_ of 500nM alone **(as already shown in Fig 4A)** however upon administering both PF-543 and Amphotericin B in combination, synergistic impact was prominent, and the IC_50_ was observed to be ∼12.5nM and 150nM for Amphotericin B and PF-543 respectively, as depicted by the isobologram micrographs in promastigotes **(Fig 7A).**

**Fig 7.**
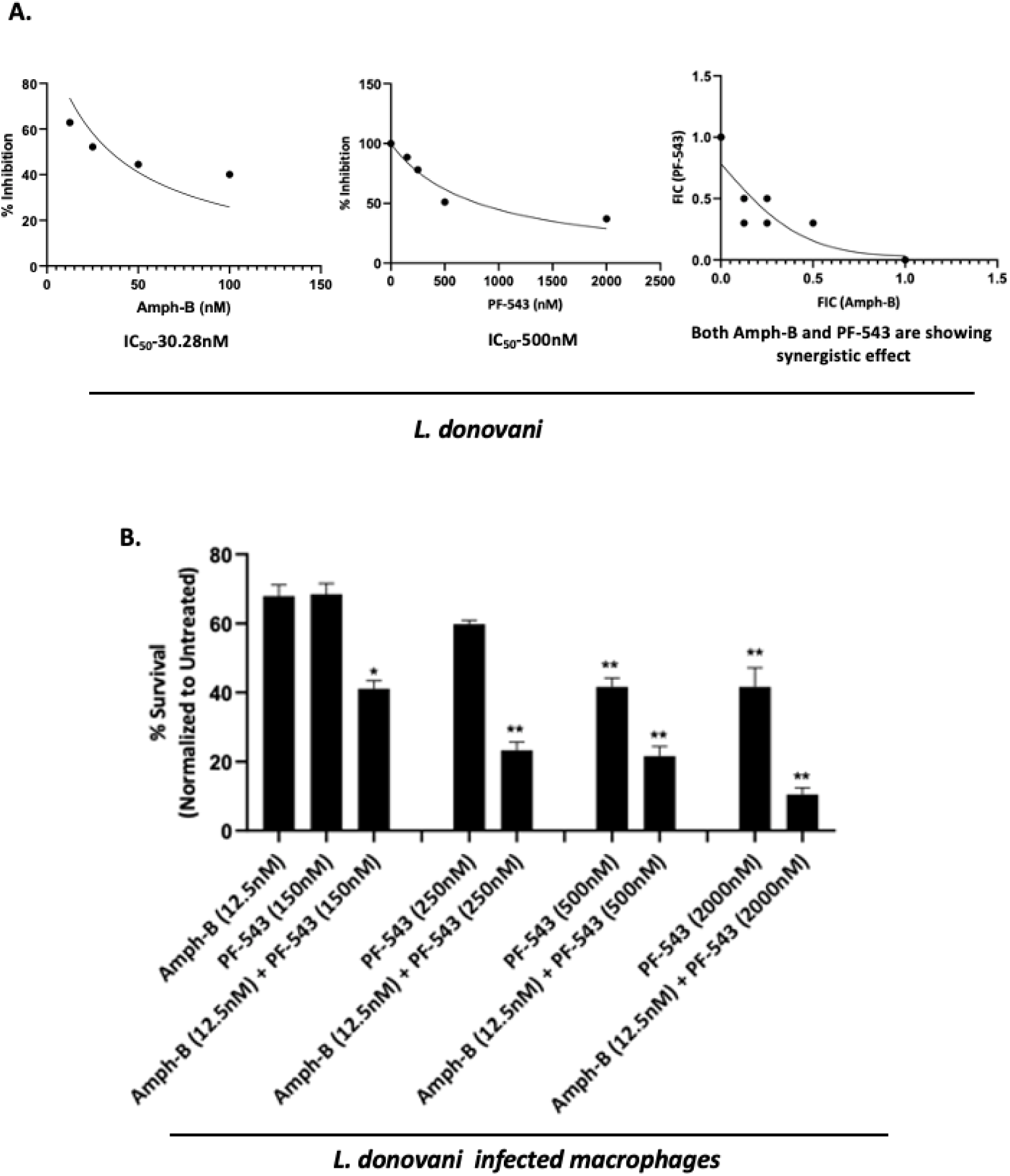
(A-B). Evaluation of PF-543 and Amphotericin B formulation on cytotoxicity or metabolic viability of *Leishmania* promastigotes and intracellular amastigotes. **(A)** To determine the combinatorial impact of PF-543 and Amphotericin B formulation, promastigotes of *L. donovani* were seeded in 96-well plates (3 × 10^6^ parasite/mL) in exponential growth phase at increasing concentrations (12.5nM–500nM) of Amphotericin B and (150nM-2000nM) of PF-543 compounds and maintained at 26°C. After 48 h of incubation, the IC_50_ was determined by MTT assay. The absorbance was measured in a MultiskanEX photometric plate reader for microplates at 540 nm. Data were obtained from three independent experiments performed in triplicate. The fractional inhibitory concentration index (FIC) was evaluated to determine the synergistic activity of lead compounds; PF-543 with Amph-B. Finally, FIC < 0.5 represented strong synergistic activity between two drugs. While FIC > 4 is was considered as the antagonistic activity. FIC values were obtained from three independent experiments. **(B)** For validating the antileishmanial activity by PF-543 and Amphotericin B formulation on intracellular amastigotes, qPCR-based analysis was performed for all the treatments. For the same, THP-1 cells were seeded at a concentration of 8 × 10^5^ cells/mL in 96-well plates and incubated for 24h with PMA (10ng/mL) supplemented with RPMI 1640. Following 24h priming with PMA, the culture medium was removed, and the cells were infected with *L. donovani* followed by treatment with the formulations having concentrations kept similar as used previously for promastigotes at 37°C and 5% CO_2_. Total RNA was extracted, and the resulting cDNA was subjected to real-time PCR analysis using primers specific for kinetoplast minicircle DNA (kDNA)/JW of *L. donovani*. This primer does not match to human genome, thus directly indicates the intracellular parasite load against the treatments. Data analysis was performed using the 2^-ΔΔCT^method.Values are mean ± S.D (n = 3). The results are representative of three independent experiments performed in triplicates.

For validating the anti-leishmanial activity by PF-543 and Amphotericin B formulation on intracellular amastigotes, qPCR-based analysis was performed for all the treatments. Here-in the synergistic treatment of Amphotericin B (12.5nM) and PF-543 (150nM) formulation to intracellular amastigotes reduced the parasite growth by 60%. Whereas, treatment with PF-543 or Amphotericin B alone decreased the parasite burden by 30%. Further reduction of 40% and 60% in parasite growth was observed with usage of higher dosage of PF-543 alone (250nM, 500nM and 2000nM), while these doses in combination with Amphotericin-B (12.5nM + 150nM, 12.5nM + 250nM, 12.5nM + 500nM, 12.5nM + 2000nM) showed higher reduction of parasite growth to ∼80-90% **(Fig 7B).** These data clearly suggested that PF-543 along with Amphotericin B can be a promising potent formulation to be tested.

### 8. Elucidation of inflammatory responses and infectivity post-PF-543 treatment in *Leishmania*-infected Swiss mice

To check the anti-leishmanial activity of PF-543 *in vivo*, the experimental group of mice were infected with *L. donovani* 24h prior to injection with 10mg/kg PF-543 and 2mg/kg amphotericin B alone via I.P route for 3 consecutive days at a dose of body weight, as a Control. While evaluating the synergistic formulation, comprising 2mg/kg PF-543 and 0.4mg/kg amphotericin B, another set of experimental mice were treated once per day for three consecutive days. Evaluation of the immunomodulatory potential was done using cytokine profiling in *Leishmania* infected Swiss mice treated with PF-543 and Amphotericin-B alone and simultaneously followed by determination of parasite infectivity using *in-vitro* parameters. To evaluate the parasitemia and expression of inflammatory markers in infected mice upon drug administration, spleen was harvested from each group of mice and qRT-PCR analysis was performed using primers specific for inflammatory markers; *mTNF-*α and *mIL-10* **(Table S1)** and parasite specific kinetoplast minicircle gene; *JW* in PF-543 and Amph-B treated infected mice in respect to the untreated Swiss mice.

In concordance with the published studies, here-in, upon analysis, treatments with either Amphotericin B or PF-543 alone in infected mice resulted in a significant decrease in *IL-10* expression, whereas increased *TNF-*α expression was observed **(Fig 8A).** Further, this effect on inflammatory markers was more pronounced upon the synergistic treatment of Amphotericin B and PF-543 formulation to *Leishmania* infected Swiss mice **(Fig 8A).**

**Fig 8.**
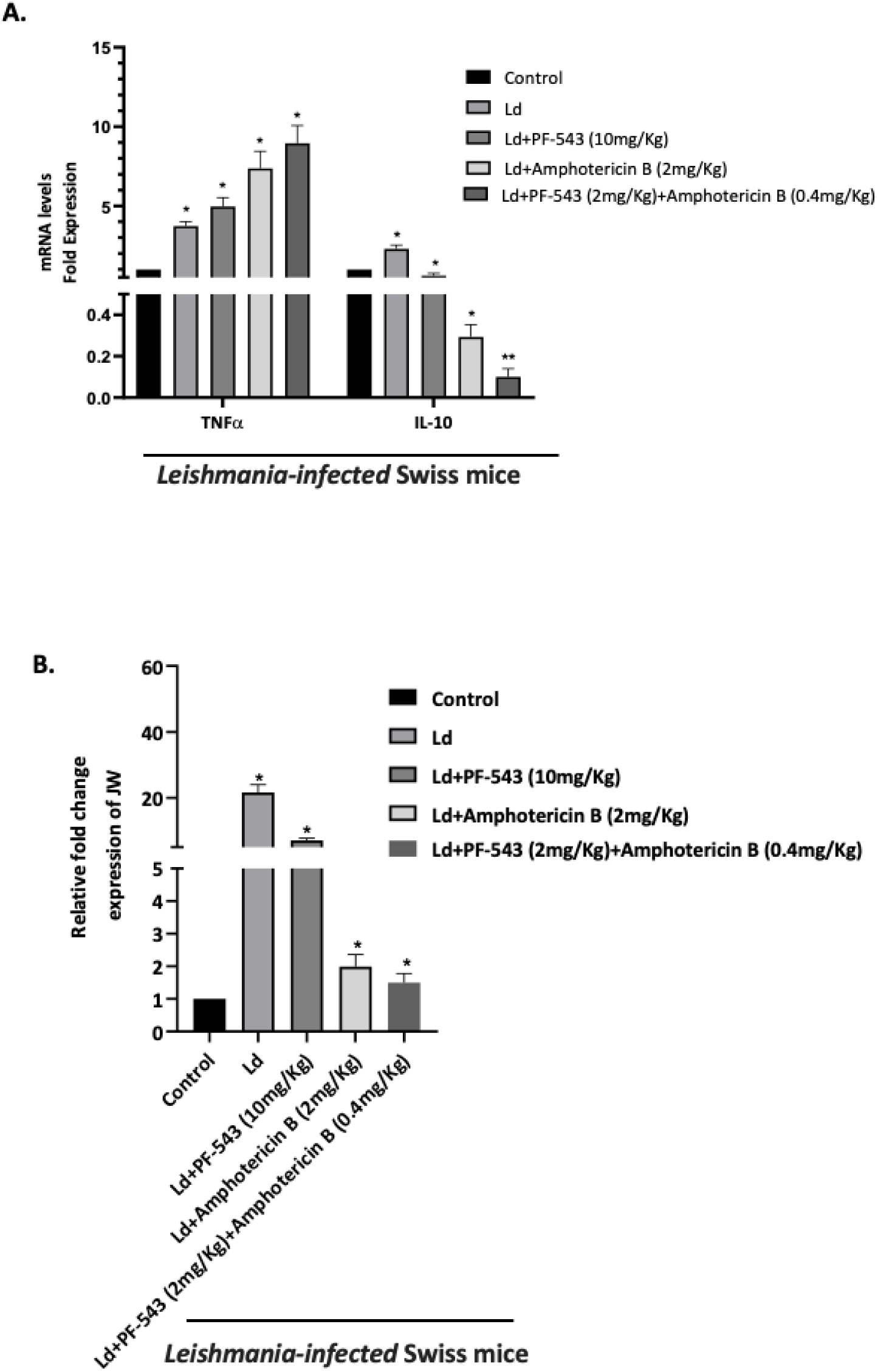
Expression of JW and inflammatory markers upon PF-543 treatment in *Leishmania-infected* Swiss mice. Evaluation of the **(A)** immunomodulatory potential and **(B)** infectivity of PF-543 and Amph-B in *Leishmania*-infected Swiss mice was performed. 10mg/Kg of PF-543 and 2mg/Kg of Amphotericin B alone were given for individual treatment in *Leishmania* infected mice. For constituting the synergistic effect of two drug partners; 2mg/Kg (1mM) of PF-543 and 0.4mg/Kg (0.12µM) of Amphotericin B were prepared in 1XPBS and given together to a mouse weighing 30gm. To evaluate the parasitemia and expression of inflammatory markers in infected mice upon drug administration, spleen was harvested from each group of mice and qRT-PCR analysis was performed using primers specific for inflammatory markers *IL-10 and TNFα* and parasite specific kinetoplast minicircle gene, *JW* in PF-543 and Amph-B treated infected mice in respect to the untreated Swiss mice. RNU6A (RNA, U6 small nuclear 1; THP-1 cells) was used as a housekeeping gene. The results were expressed as normalized fold-change of respective control. Statistical significance was quantified using the unpaired t-test. The data is a representation of mean ± SD from three independent experiments *, p< 0.05; ***, p < 0.001.

Most importantly, parasite load was also observed to be reduced upon treatment with PF-543 or Amphotericin B alone in infected mice **(Fig 8B)** where ∼33% and ∼80% reduction in parasite growth was observed respectively. Further, this down-regulation in parasite burden was drastically reduced to >90% (as demarcated by reduction in *JW* transcript) upon synergistic treatment of PF-543 and Amphotericin B formulation **(Fig 8B).**

## Discussion

The current study elucidates the role of sphingosine kinase 1 (SphK1) in the pathogenesis of *Leishmania donovani* and its potential as a therapeutic target. Our findings demonstrate that inhibition of SphK1 using PF-543, a selective inhibitor, leads to significant disruption in the parasite’s survival, proliferation, and host cell interactions. These results align with previous studies that have shown the critical role of SphK1 in various protozoan parasites, suggesting that targeting this enzyme can impair parasite viability [53,54]. Furthermore, the combination of PF-543 with Amphotericin B shows promise as a synergistic treatment strategy, with enhanced efficacy against both promastigotes and intracellular amastigotes. This is consistent with recent reports where drug combinations have been shown to enhance treatment efficacy and reduce resistance [55,56].

We successfully cloned and expressed recombinant *Leishmania* SphK1 (*rLd*SphK1) in a prokaryotic system and confirmed its enzymatic activity through fluorometric assays. Similar approaches to express and characterize SphK1 from *Leishmania* and other protozoan parasites have been reported, confirming its vital role in their metabolic pathways [57]. The results indicated that PF-543 inhibited the enzymatic activity of *rLd*SphK1 in a dose-dependent manner, reducing the levels of sphingosine-1-phosphate (S1P), a key signalling molecule that regulates cell growth and survival [58]. Additionally, the subcellular localization of *rLd*SphK1 in *Leishmania* promastigotes was primarily cytoplasmic and membranous, aligning with previous reports on SphK1 localization in other organisms [59,60].

Computational docking studies revealed that PF-543 binds effectively to the active site of *Ld*SphK1, with favourable binding interactions between key amino acid residues (e.g., ARG589, GLU586). These findings are supported by similar studies where small-molecule inhibitors have been designed to target the SphK1 enzyme, offering a potential strategy to selectively block SphK1 in parasitic infections [61]. The dissociation constant (K_D_) of 29.3µM, measured by Microscale Thermophoresis (MST), confirmed strong binding between PF-543 and *rLd*SphK1. These biophysical and computational analyses strengthen the hypothesis that PF-543 directly targets and inhibits SphK1 activity in *Leishmania*. Similar experiments were performed with human SphK1 (*rh*SphK1), and PF-543 exhibited a comparable inhibitory effect, with a slightly weaker binding affinity (K_D_ = 34.3µM for *rh*SphK1 vs. 70.6µM for SphK2) in MST assays. This suggests that PF-543 could potentially inhibit human SphK1 with similar efficiency, further justifying its consideration for therapeutic use in both parasitic and human diseases.

The study extended to *Leishmania* promastigotes with overexpressed SphK1 (*Ld*SphKa). PF-543 treatment showed a higher IC_50_ in these overexpressor parasites, suggesting that increased SphK1 expression might partially confer resistance to the inhibitor. However, even in these overexpressor strains, PF-543 still exhibited a ∼40% reduction in parasite load within infected macrophages. This emphasizes the importance of targeting SphK1 in *Leishmania* and suggests that while overexpression may confer partial resistance, inhibition of SphK1 still holds therapeutic potential, as also observed in studies targeting SphK1 in cancer cells [62].

The cytotoxic effect of PF-543 was evaluated in both *Leishmania* promastigotes and human THP-1 macrophages. The results indicated that PF-543 had a lower cytotoxic effect on THP-1 cells, with an IC_50_ of 90µM, compared to its potent action on *Leishmania* promastigotes and intracellular amastigotes. This selective toxicity aligns with findings previously reported [63], where SphK1 inhibition was shown to preferentially target pathogens without causing significant harm to host cells. The IC_50_ of 500nM highlighted PF-543’s preferential toxicity toward the parasite, reinforcing the idea that targeting SphK1 can selectively inhibit parasite growth without major toxicity to host cells.

Furthermore, PF-543 treatment led to reduced intracellular parasite burden in THP-1 cells, with a 40% reduction in parasite load, confirming its efficacy in controlling the infection. This result is consistent with similar findings in other studies, where SphK1 inhibitors significantly reduced parasite burden in macrophages infected with *Leishmania* species [64]. This was further supported by RT-PCR analysis showing decreased parasite-specific mRNA levels, indicating that PF-543 effectively reduces parasite survival within host cells.

*Leishmania* infection leads to increased ceramide generation, which disrupts membrane integrity and impairs antigen presentation, contributing to the pathogenesis [65]. Our study demonstrated that inhibition of ceramide synthesis using DL-threo-PPMP led to a decrease in parasite viability, as evidenced by increased caspase-9 activity and a decreased GSH/GSSG ratio. Interestingly, exogenous addition of sphingosine-1-phosphate (S1P) reversed the cytotoxic effect of ceramide, suggesting that S1P acts as a protective factor against ceramide-induced parasite death. These findings underscore the complex interplay between sphingolipid metabolism and parasite survival and highlight the potential therapeutic value of modulating this pathway [66].

SphK1 is known to play a crucial role in modulating the inflammatory response, and its inhibition has the potential to alter host immune responses. In *Leishmania*-infected macrophages, PF-543 treatment resulted in reducing the levels of the anti-inflammatory cytokine IL-10. This shift towards a pro-inflammatory profile is consistent with the known immunomodulatory effects of sphingosine kinase inhibition [67,68]. This may contribute to enhanced host defense against the infection, as IL-12 and TNF-α are critical for controlling *Leishmania* replication [69]. The decrease in IL-10 expression could help mitigate the immune suppression typically induced by *Leishmania*, which facilitates parasite survival.

This suggests that PF-543 not only targets the parasite directly but also modulates the host immune environment to favour parasite clearance.

As per previous reports, Amphotericin B, a widely used anti-leishmanial drug, targets the parasitic membrane by binding to ergosterol, but its use is limited due to toxicity and resistance concerns [70]. Our study explored the combination of PF-543 and Amphotericin B, finding a synergistic effect. The IC_50_ values for the combination treatment were significantly lower compared to individual drugs, with a greater than 80% reduction in parasite burden in both promastigotes and intracellular amastigotes. This synergy may be due to PF-543’s ability to alter sphingolipid metabolism, affecting membrane integrity and facilitating increased susceptibility to Amphotericin B-induced damage, as demonstrated in previous work by Al-Rawi *et al* [71].

Further, *in vivo* studies in *Leishmania*-infected Swiss mice showed that PF-543, either alone or in combination with Amphotericin B, significantly reduced the parasite load in spleen tissues. Treatment with the combination therapy resulted in a >90% reduction in parasite burden, compared to a ∼33% reduction with PF-543 alone and ∼80% reduction with Amphotericin B alone. Cytokine analysis revealed an increase in TNF-α expression and a decrease in IL-10, further supporting the immune-modulatory role of PF-543 *in vivo*. These findings are consistent with studies showing that combination therapies enhance anti-parasitic efficacy and improve immune responses in *Leishmania*-infected models [72].

In conclusion, this study provides compelling evidence for the crucial role of SphK1 in *Leishmania* survival and pathogenesis and highlights the potential of PF-543 as a selective inhibitor. The combination of PF-543 with Amphotericin B shows significant promise as a synergistic treatment strategy against *Leishmania donovani*. Further investigation into the long-term effects and potential clinical applications of this combination therapy is warranted,particularly in the context of combating drug resistance and enhancing immune responses in infected hosts. Additionally, modulating sphingolipid metabolism offers a novel therapeutic approach for leishmaniasis.

## Acknowledgments

Singh S conceptualized the work, designed the experiments, interpreted the data and edited the manuscript. EM executed the experiments, performed the data interpretation and wrote the manuscript. EM, RB, JS, MS, SS and NJ performed immunological studies, microscopy, molecular assays, parasite culture, cloning, protein purification and mice experiments. This study was funded by CSIR Scientist Pool Officer (SRA) funding agency research grant (13(9160-A)2021-Pool) provided to Dr. Evanka Madan. RB and JS were supported by DHR-Women Scientist and BioCARe Women Scientist from DBT respectively. SS is a recipient of the National Bio Scientist Award. All authors contributed to the article and approved the submitted version. The work in Shailja Singh’s lab at Special Centre for Molecular Medicine (SCMM), JNU, is supported by IRHPA IPA/2020/000007, Department of Science and Technology and Drug and Pharmaceuticals Research Program (DPRP) (Project No. P/569/2016-1/TDT, SS). We would like to acknowledge Prof. Md. Imtiyaz Hassan (Professor, Structural Biology Laboratory, Jamia Millia Islamia) for providing us with the Human SphK1 clone and Prof. Zhang, Kai (Professor, Department of Biological Sciences, Texas Tech University) for providing us with pXG-GFP-SKa plasmid which we used to overexpress SKa (or SphK1) in *L. donovani*. The funders had no role in the study design; collection, analysis, and interpretation of data; in the writing of the report; and in the decision to submit this article for publication.

## Plagiarism Check

This manuscript has been screened for plagiarism and found to be original.

---

**Fig S1.**
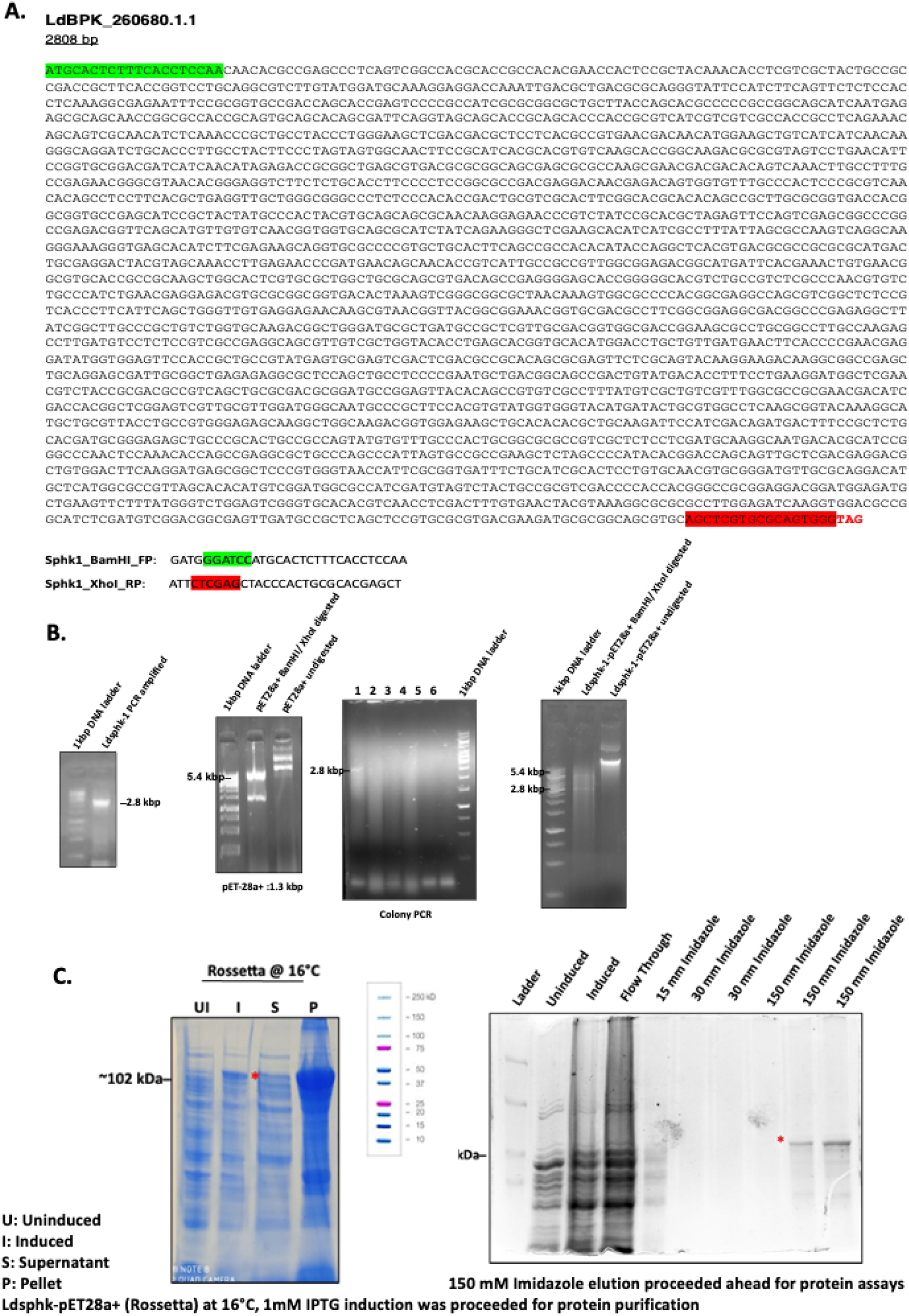
(A) Nucleotide sequence encoding full length sequence of *L. donovani* Sphk1 (1-2808 bp). This sequence was selected for recombinant protein generation as a fusion protein with an N-terminal 6x-Histidine-tag using the vector pET-28 a(+) vector. **(B) Cloning of *L. donovani* SphK1 in pET28a(+) vector.** PCR amplification of DNA fragment from genomic DNA of *L. donovani* using *Ld*Sphk1_BamHI and *Ld*Sphk1_XhoI with Phusion™ High-Fidelity DNA Polymerase. Purification of the amplified DNA fragment was done using QIAquick Gel Extraction Kit followed by digestion of purified *Ld*Sphk1 insert and the expression vector pET-28 a(+) with BamHI/XhoI restriction enzymes. The positive clones were screened by colony PCR followed by confirmation of the cloned plasmid with BamHI/XhoI restriction digestion. **(C) *rLd*Sphk1 recombinant protein purification**. Transformation of positive clone in Rossetta *E. coli* expression strain for *Ld*Sphk1 recombinant protein expression. For recombinant protein induction, the cells were induced with 1mM IPTG for 4 hours at 37°C in Terrific broth. Purification of recombinant protein was achieved by affinity chromatography. The *rLd*SphK1 protein was eluted after addition of 150mM imidazole solution. The eluted fractions were analysed by running on 12% SDS PAGE.

**Fig S2.**
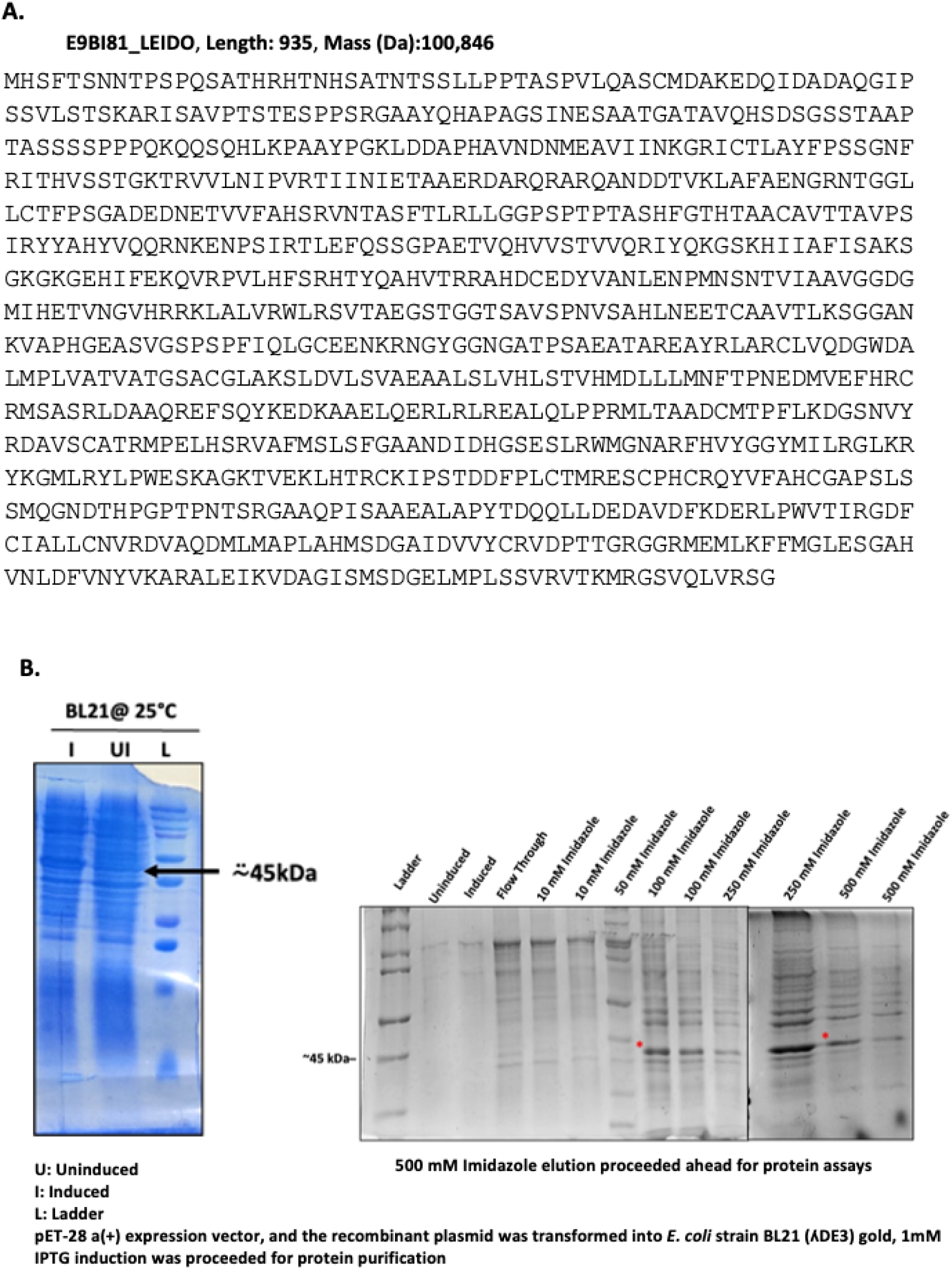
(A) The Human SphK1 amino acid sequence for the protein. (B) Purification of the recombinant *Human*SphK1 (*rHuman*SphK1) protein. Overexpression of 6×His-SphK-1 (*rhSphK-1*) was induced with 1mM IPTG at an optical density (OD600) of 0.6, for 4 h at 37°C. The protein was purified using Ni-NTA agarose resin. The *rhSphK1* protein was eluted with a continuous imidazole gradient of 50mM, 100mM, 250mM, and 500mM. The protein purification was validated by 12% SDS-PAGE, followed by immunoblotting with anti-His tag antibody. A single band corresponding to ∼50 kDa was observed on SDS PAGE corresponding to Human specific SphK1.

**Fig S3.**
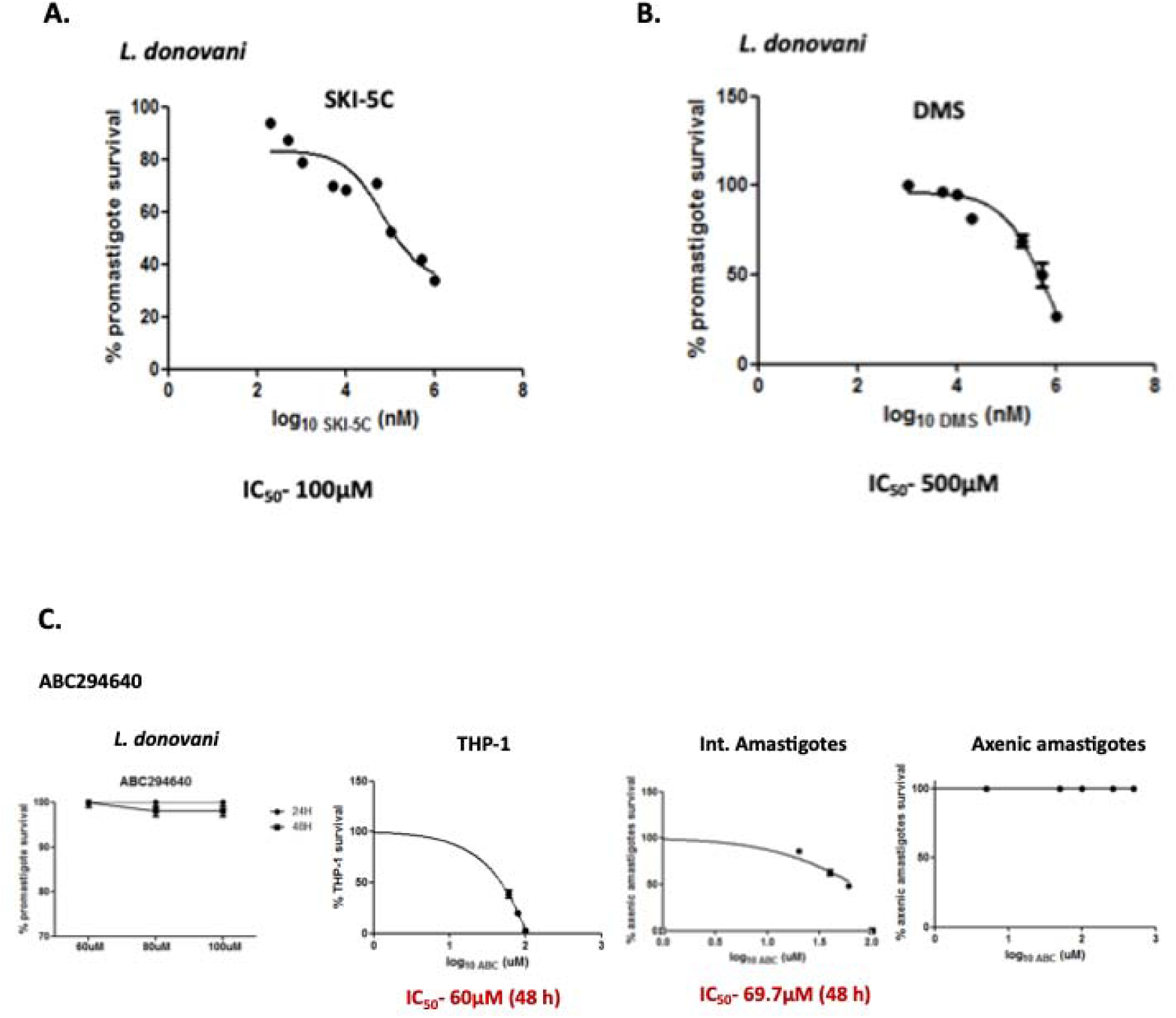
(A,B) Effect of SphK1 inhibitors; SKI-5C and DMS in *Leishmania promastigotes*. To evaluate the IC_50_ for SKI-5C and DMS, approximately 5×10^4^ *Ld* Bob cells were seeded in each well of 96-well flat bottom plates and supplemented with M199 media containing SKI-5C and DMS (200µl/well) respectively. The cells were further incubated for 2 days at 37°C and 5% CO_2_. The IC_50_ were found to be 100μM for SKI-5C and 500μM for DMS respectively in *Leishmania donovani* promastigotes. Each experiment was done in triplicates and repeated thrice. **(C) Effect of SphK1 inhibitor; ABC294640 in *Leishmania spp* and host macrophages.** To evaluate the IC_50_ for ABC294640, approximately 6×10^3^ THP-1 and 5×10^4^ *Ld* Bob cells were seeded in each well of 96-well flat bottom plates and supplemented with RPMI and M199 media containing ABC294640 (200µl/well) respectively. For intracellular amastigotes, 1 × 10^6^ THP-1 cells, treated with 50 ng/ml of phorbol 12-myristate 13-acetate (PMA) were seeded on glass coverslip in a 6-well plate for 48h. They were infected with late log-phase *L. donovani* promastigotes and simultaneously treated with ABC294640. The cells were further incubated for 2 days at 37°C and 5% CO_2_. To determine, the intracellular parasite burdens (mean number of amastigotes per macrophage) were microscopically assessed using Giemsa staining. For axenic amastigotes, the axenically cultured forms grew optimally at a temperature of 32-33°C in a growth media with pH of 5.4. The IC_50_ were found to be 60μM for THP-1 macrophages and 69.7μM for intracellular amastigotes respectively. Each experiment was done in triplicates and repeated thrice.

**Fig S4.**
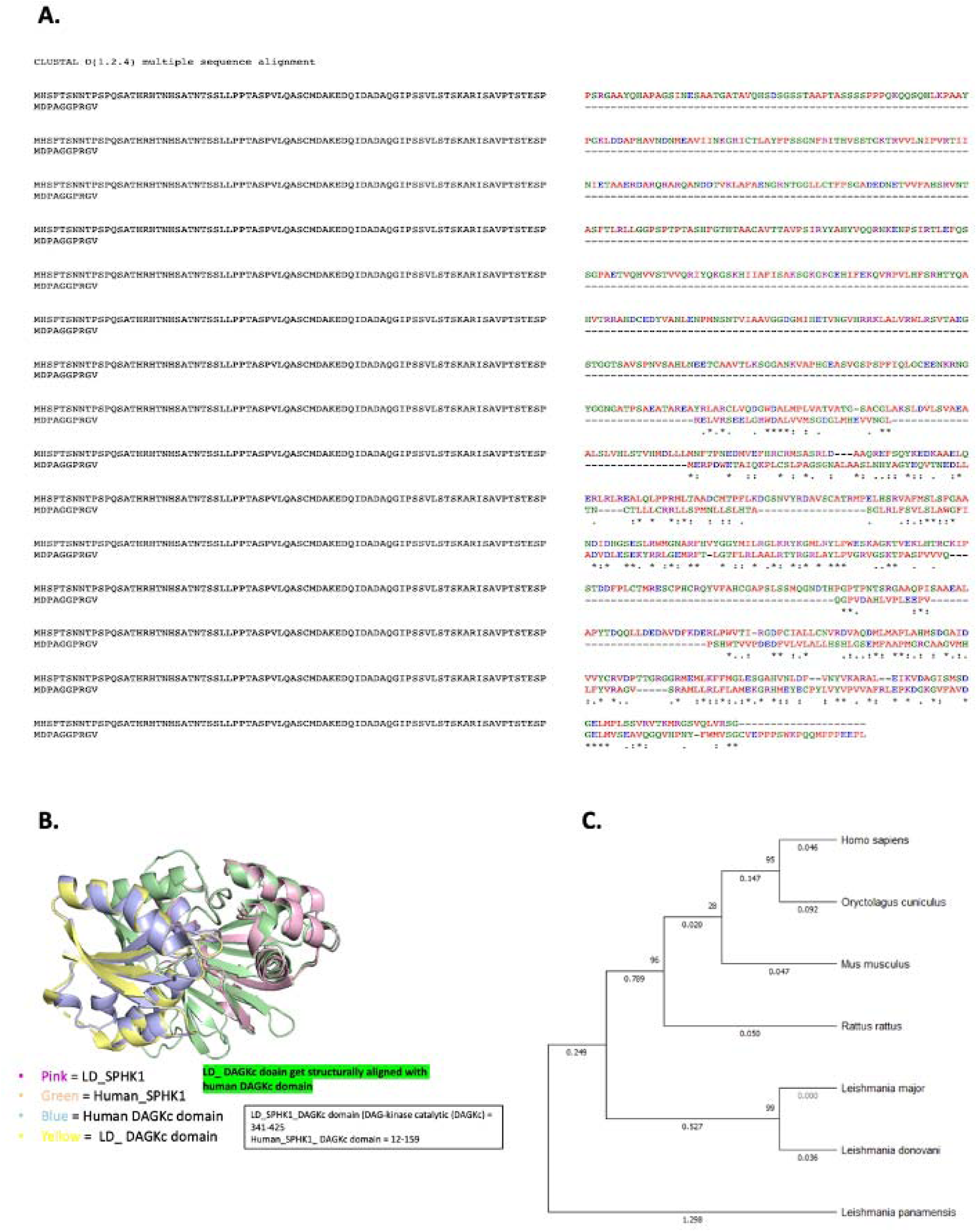
(A) Amino acid sequence alignment of LD_SphK1 and Human_SphK1 utilizing MEGAX program (version 10.2.2). The results demonstrated considerable alignment with similar and a limited number of identical amino acids. **(B) 3D structural alignment of LD_SphK1 and Human_SphK1 protein structures with Pymol software.** Substantial alignment between the two proteins, with an RMSD of 0.585 was observed. Furthermore, the alpha helix and beta sheets of the Human DAGKc domain (Blue) accurately aligned with the LD_DAGKc domain (Yellow). (C) Construction of phylogenetic tree utilizing SphK1 gene sequences from various species of *Leishmania,* humans, rats, and mice. The phylogenetic tree was constructed using neighbor-joining technique with MEGAX software. Bootstrap analyses with 1000 replicates were conducted to evaluate the robustness of the constructed phylogenetic tree.

**Fig S5.**
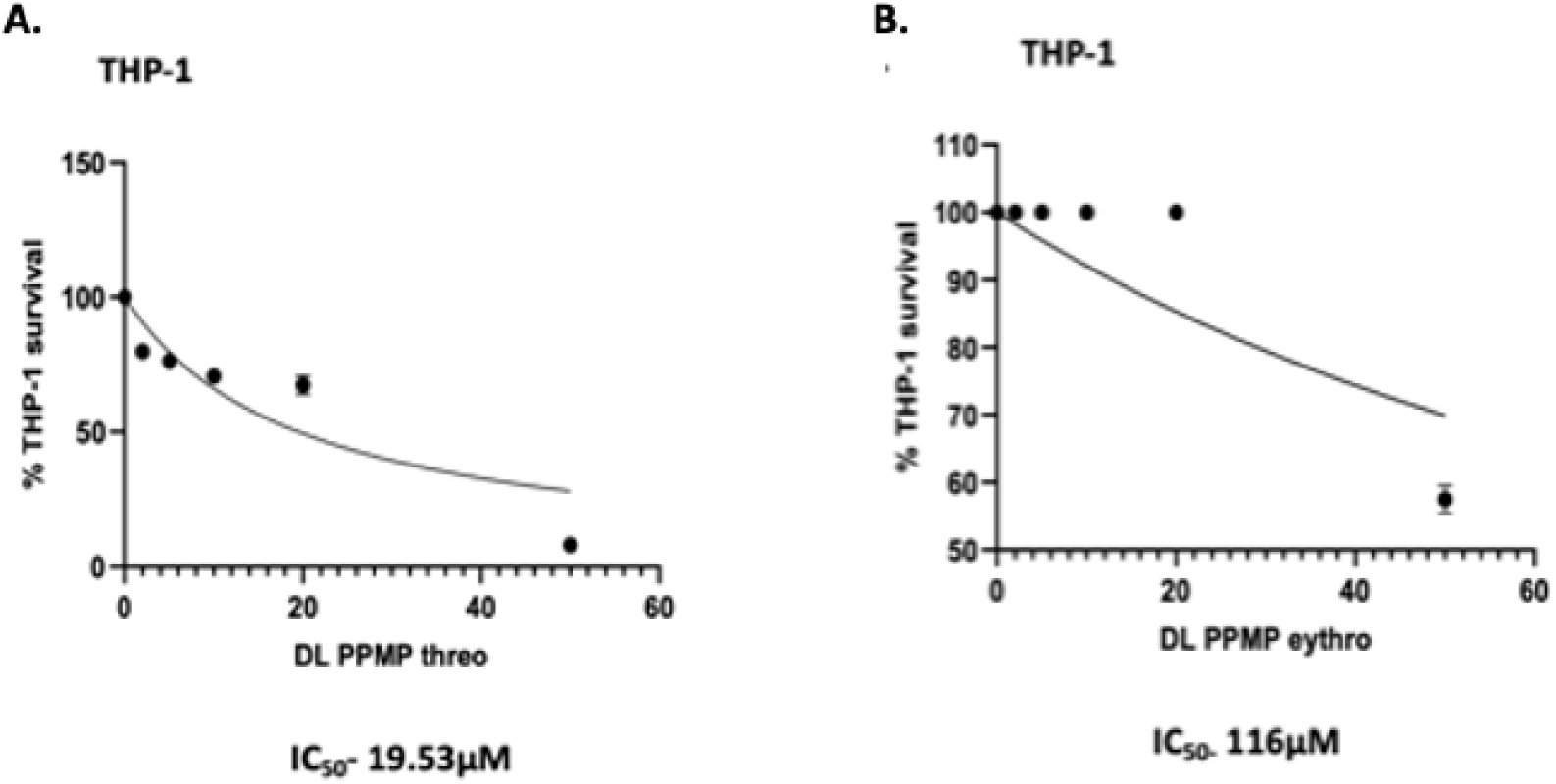
(A) Effect of Ceramide analogs; DL-threo-PPMP and DL-erythro-PPMP in THP-1 macrophages. To evaluate the IC_50_ for DL-threo-PPMP and DL-erythro-PPMP, approximately 6×10^3^ THP-1 cells were seeded in each well of 96-well flat bottom plates and supplemented with RPMI media containing DL-threo-PPMP and DL-erythro-PPMP (200µl/well) respectively. The cells were further incubated for 2 days at 37°C and 5% CO_2_. The IC_50_ were found to be 19.53μM and 116μM for DL-threo-PPMP and DL-erythro-PPMP respectively in THP-1 macrophages. Each experiment was done in triplicates and repeated thrice.

**Table S1.**
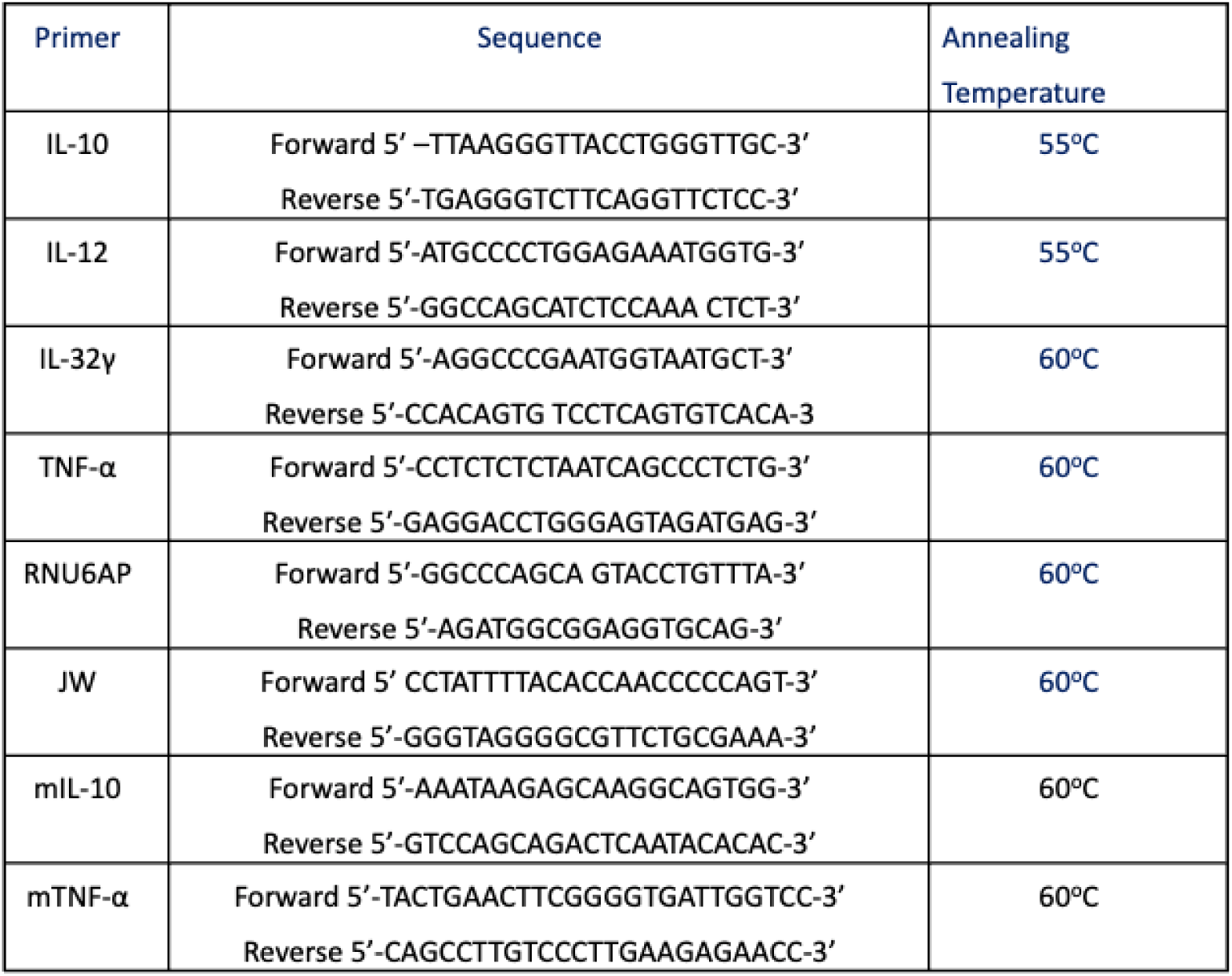
Primer sequences of target genes mentioned along with their annealing temperatures.

## References

1. Alvar J, Vélez ID, Bern C, Herrero M, Desjeux P, Cano J, et al. Leishmaniasis worldwide and global estimates of its incidence. PLoS One. 2012;7:e35671.

2. Arango Duque G, Descoteaux A. Leishmania survival in the macrophage: Where the ends justify the means. Curr Opin Microbiol. 2015;26:32–40.

3. Rub A, Arish M, Husain SA, Ahmed N, Akhter Y. Host-lipidome as a potential target of protozoan parasites. Microbes Infect. 2013;15:649–60.

4. Bhargava P, Singh R. Developments in diagnosis and antileishmanial drugs. Interdiscip Perspect Infect Dis. 2012;2012:626838.

5. Mahmoudvand H, Sharififar F, Sharifi I, Ezatpour B, Fasihi Harandi M, Makki MS, et al. In vitro inhibitory effect of Berberis vulgaris and its main component, berberine against different Leishmania species. Iran J Parasitol. 2014;9:28–36.

6. Moore EM, Lockwood DN. Treatment of visceral leishmaniasis. J Glob Infect Dis. 2010;2(2):151–8.

7. Datta A, Podder I, Das A, Sil A, Das NK. Therapeutic modalities in post kala-azar dermal leishmaniasis: A systematic review of the effectiveness and safety of the treatment options. Indian J Dermatol. 2021;66(1):34–43.

8. Huang LS, Berdyshev E, Mathew B, Fu P, Gorshkova IA, He D, et al. Targeting sphingosine kinase 1 attenuates bleomycin-induced pulmonary fibrosis. FASEB J. 2013;27:1749–60.

9. Chen J, Tang H, Sysol JR, Moreno-Vinasco L, Shioura KM, Chen T, et al. The sphingosine kinase 1/sphingosine-1-phosphate pathway in pulmonary arterial hypertension. Am J Respir Crit Care Med. 2014;190:1032–43.

10. Price MM, Oskeritzian CA, Falanga YT, Harikumar KB, Allegood JC, Alvarez SE, et al. A specific sphingosine kinase 1 inhibitor attenuates airway hyperresponsiveness and inflammation in a mast cell-dependent murine model of allergic asthma. J Allergy Clin Immunol. 2013;131:501–11.e1.

11. Wang L, Chen F, Pan Y, Lin L, Xiong X. Effects of FTY720 on lung injury induced by hindlimb ischemia reperf2usion in rats. Mediators Inflamm. 2017;2017:5301312.

12. Nishiuma T, Nishimura Y, Okada T, Kuramoto E, Kotani Y, Jahangeer S, et al. Inhalation of sphingosine kinase inhibitor attenuates airway inflammation in asthmatic mouse model. Am J Physiol Lung Cell Mol Physiol. 2008;294:L1085–93.

13. Arish M, Husein A, Ali R, Tabrez S, Naz F, Ahmad MZ, et al. Sphingosine-1-phosphate signaling in Leishmania donovani infection in macrophages. PLoS Negl Trop Dis. 2018;12:e0006360.

14. Pulkoski-Gross MJ, Obeid LM. Molecular mechanisms of regulation of sphingosine kinase 1. Biochim Biophys Acta Mol Cell Biol Lipids. 2018;1863:1413–22.

15. Zhang O, Hsu FF, Xu W, Pawlowic M, Zhang K. Sphingosine kinase A is a pleiotropic and essential enzyme for Leishmania survival and virulence. Mol Microbiol. 2013;90(3):489–501.

16. Cao M, Ji C, Zhou Y, Huang W, Ni W, Tong X, et al. Sphingosine kinase inhibitors: A patent review. Int J Mol Med. 2018;41:2450–60.

17. Patel DS, Ahmad F, Abu Sneineh M, Patel RS, Reddy SR, Llukmani A, et al. The importance of sphingosine kinase in breast cancer: A potential for breast cancer management. Cureus. 2021;13(2):e13413.

18. Kim SB, Limbu KR, Oh YS, Kim SL, Park SK, Baek DJ, et al. Novel dimer derivatives of PF-543 as potential antitumor agents for the treatment of non-small cell lung cancer. Pharmaceutics. 2022;14(2):369.

19. Yakhak Hoeji. Antitumor activity of PF-543 and PF-543 derivative (22c) as sphingosine kinase 1 inhibitors. Yakhak Hoeji. 2018;62(5):293–6.

20. Sah RK, Anand S, Dar W, Jain R, Kumari G, Madan E, et al. Host-erythrocytic sphingosine-1-phosphate regulates Plasmodium histone deacetylase activity and exhibits epigenetic control over cell death and differentiation. Microbiol Spectr. 2023;11(2):e02766–22.

21. Romano PS, Akematsu T, Besteiro S, Bindschedler A, Carruthers VB, Chahine Z, et al. Autophagy in protists and their hosts: When, how and why? Autophagy Reports. 2023;2(1):2149211.

22. New RR, Chance ML, Heath S. Antileishmanial activity of amphotericin and other antifungal agents entrapped in liposomes. J Antimicrob Chemother. 1981;7(4):371–81.

23. Verma J, Verma L, Tripathi K. Sterol enriched mixed lamellarity amphotericin intercalating liposomes in saline and the process for their preparation. US Patent 2007/0218119 A1.

24. Aslett M, Aurrecoechea C, Berriman M, Brestelli J, Brunk BP, Carrington M, et al. TriTrypDB: a functional genomic resource for the Trypanosomatidae. Nucleic Acids Res [Internet]. 2010 [cited 2023 Dec 21];38(Database issue):D457–62.

25. Troshin PV, Procter JB, Barton GJ. Java bioinformatics analysis web services for multiple sequence alignment—JABAWS:MSA. Bioinformatics. 2011;27(14):2001–2.

26. Kumar S, Stecher G, Li M, Knyaz C, Tamura K. MEGA X: Molecular evolutionary genetics analysis across computing platforms. Mol Biol Evol. 2018;35(6):1547–9.

27. DeLano WL. PyMOL: An open-source molecular graphics tool. CCP4 Newsl Protein Crystallogr [Internet]. 2002;40:82–92.

28. Berman HM. The Protein Data Bank. Nucleic Acids Res. 2000;28(1):235–42.

29. The UniProt Consortium. UniProt: a hub for protein information. Nucleic Acids Res. 2015;43(Database issue):D204–12.

30. Ivens AC, Peacock CS, Worthey EA, Murphy L, Aggarwal G, Berriman M, et al. The genome of the kinetoplastid parasite, Leishmania major. Science. 2005;309(5733):436–42.

31. Kelley LA, Mezulis S, Yates CM, Wass MN, Sternberg MJ. The Phyre2 web portal for protein modeling, prediction and analysis. Nat Protoc [Internet]. 2015 [cited 2021 Aug 7];10(6):845–58. Available from: https://www.nature.com/articles/nprot.2015.053

32. Laskowski RA, MacArthur MW, Moss DS, Thornton JM. PROCHECK: a program to check the stereochemical quality of protein structures. J Appl Crystallogr. 1993;26(2):283– 91.

33. Kaplan W, Littlejohn TG. Swiss-PDB Viewer (DeepView): An open-source molecular modeling program. Brief Bioinform. 2001;2(2):195–7.

34. Milne GWA. Computer software review of ChemBioDraw 12.0. J Chem Inf Model. 2010;50(11):2053.

35. Morris GM, Lim-Wilby M. Molecular docking. Methods Mol Biol. 2008;443:365–82.

36. Racine J. The Cygwin tools: a GNU toolkit for Windows. J Appl Econom [Internet]. 2000;15(3):331–41.

37. Salentin S, Schreiber S, Haupt VJ, Adasme MF, Schroeder M. PLIP: fully automated protein–ligand interaction profiler. Nucleic Acids Res [Internet]. 2015;43(W1):W443–7.

38. Laskowski RA, Swindells MB. LigPlot+: multiple ligand-protein interaction diagrams for drug discovery. J Chem Inf Model. 2011;51(10):2778–86.

39. Miyata T. Discovery Studio modeling environment. Ensemble. 2015;17:98–104.

40. DeLano WL. PyMOL: An open-source molecular graphics tool. CCP4 Newsl Protein Crystallogr [Internet]. 2002;40:82–92.

41. Vallur AC, Tutterrow YL, Mohamath R, Pattabhi S, Hailu A, Abdoun AO, et al. Development and comparative evaluation of two antigen detection tests for visceral leishmaniasis. BMC Infect Dis. 2015;15:384.

42. Kapler GM, Coburn CM, Beverley SM. Stable transfection of the human parasite Leishmania major delineates a 30-kilobase region sufficient for extrachromosomal replication and expression. Mol Cell Biol. 1990;10(3):1084–94.

43. Pawar H, Puri M, Fischer-Weinberger R, Madhubala R, Zilberstein D. The arginine sensing and transport binding sites are distinct in the human pathogen Leishmania. PLoS Negl Trop Dis. 2019;13(1):e0007304.

44. Darlyuk I, Goldman A, Roberts SC, Ullman B, Rentsch D, Zilberstein D. Arginine homeostasis and transport in the human pathogen Leishmania donovani. J Biol Chem. 2009;284(30):19800–7.

45. Fivelman QL, Adagu IS, Warhurst DC. Modified fixed-ratio isobologram method for studying in vitro interactions between atovaquone and proguanil or dihydroartemisinin against drug-resistant strains of Plasmodium falciparum. Antimicrob Agents Chemother. 2004;48(11):4097–102.

46. Akoachere M, Buchholz K, Fischer E, Burhenne J, Haefeli WE, Schirmer RH, et al. In vitro assessment of methylene blue on chloroquine-sensitive and -resistant Plasmodium falciparum strains reveals synergistic action with artemisinins. Antimicrob Agents Chemother. 2005;49(11):4592–7.

47. Kelly JX, Smilkstein MJ, Cooper RA, Lane KD, Johnson RA, Janowsky A, et al. Design, synthesis, and evaluation of 10-N-substituted acridones as novel chemosensitizers in Plasmodium falciparum. Antimicrob Agents Chemother. 2007;51(12):4133–40.

48. Hissin PJ, Hilf R. A fluorometric method for determination of oxidized and reduced glutathione in tissues. Anal Biochem. 1976;74:214–26.

49. Rub A, Dey R, Jadhav M, Kamat R, Chakkaramakkil S, Majumdar S, et al. Cholesterol depletion associated with Leishmania major infection alters macrophage CD40 signalosome composition and effector function. Nat Immunol. 2009;10(3):273–80.

50. Lapara NJ 3rd, Kelly BL. Suppression of LPS-induced inflammatory responses in macrophages infected with Leishmania. J Inflamm (Lond). 2010;7:8.

51. Kumari S, Kumar V, Tiwari RK, Ravidas V, Pandey K, Kumar A. Amphotericin B: a drug of choice for visceral leishmaniasis. Acta Trop. 2022;235:106661.

52. Laniado-Laborín R, Cabrales-Vargas MN. Amphotericin B: side effects and toxicity. Rev Iberoam Micol. 2009;26(4):223–7.

53. Wang X, et al. Sphingosine kinase 1: a potential therapeutic target for parasitic diseases. J Parasitol Res. 2011;2011:1–10.

54. Cahill JL, et al. Sphingosine kinase 1 plays an essential role in the growth and viability of Toxoplasma gondii. Int J Parasitol. 2009;39(9):1061–70.

55. Müller MA, et al. Synergistic effects of drug combinations for the treatment of Leishmania infection. Antimicrob Agents Chemother. 2018;62(11):e01455–18.

56. Siqueira-Neto JL, et al. Sphingosine kinase inhibitors as promising new anti-leishmanial agents: synergy with other drugs. J Antimicrob Chemother. 2014;69(3):667–74.

57. Gordon JR, et al. Characterization of sphingosine kinase 1 from Leishmania donovani and its potential as a drug target. Mol Biochem Parasitol. 2012;183(1):1–9.

58. Pitson SM, et al. PF-543, a potent sphingosine kinase 1 inhibitor, reduces sphingosine-1-phosphate production and blocks the growth of T-cell acute lymphoblastic leukemia cells. Mol Cancer Ther. 2008;7(6):1559–68.

59. Bresnick AR, et al. Subcellular localization of sphingosine kinase 1 in Leishmania spp. and its association with the parasite’s membrane-bound structures. J Cell Sci. 2009;122(20):3732–42.

60. Dunn TM, et al. The subcellular localization of sphingosine kinase 1 in mammalian cells: implications for its function in cell signaling. J Lipid Res. 2010;51(10):3061–71.

61. Barauskas J, et al. Design and synthesis of small-molecule inhibitors of sphingosine kinase 1: a computational approach. Bioorg Med Chem. 2014;22(24):6625–35.

62. Kharel Y, et al. Overexpression of sphingosine kinase 1 confers partial resistance to targeted therapy in cancer cells. Mol Cancer Ther. 2016;15(12):3084–93.

63. Foster LJ, et al. Sphingosine kinase inhibition: potential for selective targeting of microbial pathogens in humans. Trends Pharmacol Sci. 2007;28(3):178–85.

64. Hernández-Ruiz J, et al. Inhibition of sphingosine kinase 1 reduces Leishmania infection in macrophages by targeting parasite survival pathways. Parasite Immunol. 2010;32(11):730– 8.

65. Liu Y, et al. Ceramide generation in Leishmania: a potential target for therapeutic intervention. J Lipid Res. 2012;53(10):2104–14.

66. Maceyka M, et al. Sphingosine kinase 1 and 2 as regulators of sphingolipid metabolism and biological function. J Lipid Res. 2012;53(8):1397–409.

67. Rennie DE, et al. Sphingosine kinase 1 inhibition modulates cytokine production in T-cell immune responses. J Immunol. 2015;194(3):1261–70.

68. Saviola AJ, et al. Sphingosine kinase 1 inhibition alters immune responses and enhances anti-parasitic immunity. PLoS Pathog. 2013;9(5):e1003351.

69. Olivier M, et al. The role of tumor necrosis factor-alpha and interleukin-12 in host resistance to Leishmania. Infect Immun. 2002;70(5):2771–5.

70. Müller M, et al. Amphotericin B for the treatment of leishmaniasis: progress and limitations. Clin Infect Dis. 2015;60(2):217–28.

71. Al-Rawi MA, et al. Synergistic effects of PF-543 with amphotericin B against Leishmania donovani: targeting sphingolipid metabolism and membrane integrity. Antimicrob Agents Chemother. 2019;63(5):e02211–18.

72. Soliman MM, et al. Combination therapy for leishmaniasis: enhanced therapeutic efficacy in Leishmania donovani-infected mouse models. Antimicrob Agents Chemother. 2014;58(10):5955–62.

